# Oligogenic rare variant contributions in schizophrenia and their convergence with genes harbouring *de novo* mutations in schizophrenia, autism and intellectual disability: Evidence from multiplex families

**DOI:** 10.1101/829101

**Authors:** Jibin John, Prachi Kukshal, Triptish Bhatia, Ricardo Harripaul, V L Nimgaonkar, S N Deshpande, B.K. Thelma

## Abstract

Clinical and genetic heterogeneity has been documented extensively in schizophrenia, a common behavioural disorder with heritability estimates of about 80%. Common and rare *de novo* variant based studies have provided notable evidence for the likely involvement of a range of pathways including glutamatergic, synaptic signalling and neurodevelopment. To complement these studies, we sequenced exomes of 11 multimember affected schizophrenia families from India. Variant prioritisation performed based on their rarity (MAF <0.01), shared presence among the affected individuals in the respective families and predicted deleterious nature, yielded a total of 785 inherited rare protein sequence altering variants in 743 genes among the 11 families. These showed an enrichment of genes involved in the extracellular matrix and cytoskeleton components, synaptic and neuron related ontologies and neurodevelopmental pathways, consistent with major etiological hypotheses. We also noted an overrepresentation of genes from previously reported gene sets with *de novo* protein sequence altering variants in schizophrenia, autism, intellectual disability; FMRP target and loss of function intolerant genes. Furthermore, a minimum of five genes known to manifest behavioural/neurological and nervous system abnormalities in rodent models had deleterious variants in them shared among all affected individuals in each of the families. Majority of such variants segregated within and not across families providing strong suggestive evidence for the genetically heterogeneous nature of disease. More importantly, study findings unequivocally support the classical paradigm of cumulative contribution of multiple genes, notably with an apparent threshold effect for disease manifestation and offer a likely explanation for the unclear mode of inheritance in familial schizophrenia.

## Introduction

Substantial but not exclusive contribution of genetic factors to the development of Schizophrenia (SZ) has been noted by classical twin, family, and adoption studies(Sullivan et al., 2003), further augmented by the high heritability estimates (up to 80%) based on genetic epidemiological studies (Cannon et al., 1998; Lichtenstein et al., 2009). Till recently, genetic advances have been made to resolve this complexity by largely relying on the common disease–common variant (CDCV) hypothesis and over 100 independent risk loci have been identified in SZ through genome wide association studies (GWAS) and their meta-analyses (Ripke et al., 2014). Further analysis showed that more than one common risk allele is present in every 1Mb genomic region across ≥71% of the total genome(Loh et al., 2015). Two other GWASs with meta-analyses have now added another 50 (Pardiñas et al., 2018) and 30 (Li et al., 2017) loci to this list. Despite such significant progress, common variant has not been able to explain the high heritability in SZ and most of the variants identified had small to modest effect sizes. On the other hand, based only on theoretical grounds an important role for rare variants in complex diseases was proposed (Pritchard, 2001). Of note, deep sequencing technologies enabled identification of such rare variants and ∼1000 rare *de novo* variants have been discovered in SZ patients, in a predominantly parent–child trio design(Li et al., 2016). Most of these variants are clustered in genes of disease related pathways such as activity-regulated cytoskeleton-associated protein (ARC), fragile-X mental retardation protein (FMRP) interacting genes, N-methyl-D-aspartate receptor (NMDA) complex, postsynaptic density (PSD)-95 complexes, calcium channels and chromatin remodelling genes(Flint, 2016; Fromer et al., 2014; McCarthy et al., 2014). However, due to their *de novo* origin, their contribution to disease heritability has not been substantiated. Involvement of immunological/infection related genes in SZ etiology has also been widely discussed from time to time (Khandaker et al., 2015; Muller and J. Schwarz, 2010). Association studies indeed identified a few loci/genes such as human leukocyteantigen (HLA), complement component 4 (C4) genes etc. (Harrison, 2015; Sekar et al., 2016), but enrichment of such genes have not been reported to date. However, a polygenic burden of rare variants has been reported in recent case-control based studies(Genovese et al., 2016; Purcell et al., 2014; Richards et al., 2016). These findings have strongly hinted at the likely involvement of genes involved in synaptic function, neurodevelopmental processes, glutamatergic signalling etc.

On the other hand, familial aggregation of SZ is reported globally with an elevated risk among the relatives of the affected individuals and the risk being directly proportional to the biological relationship with the affected individuals over that of the population at large (Chou et al., 2017). A few studies in singleton and multiplex SZ families have shown rare but highly penetrant inherited risk variants that could contribute to genetic liability. Balanced translocation t(1:11) (q43,q21) segregating with SZ and related psychiatric disorders in a large family (St Clair et al., 1990);deletion at *SLC1A1* co-segregating with bipolar/schizoaffective disorder and SZ in a five generation family (Myles-Worsley et al., 2013) are some examples. More recent whole exome sequencing (WES)/whole genome sequencing in small/modest sized multiplex families have added to the list of inherited rare protein coding variants. These include variants in previously analyzed candidate genes namely *RELN*(Zhou et al., 2016), *UNC13B*(Egawa et al., 2016), *GRM5, LRP1B, and PPEF2* (Timms et al., 2013),*GRIN3B* (Hornig et al., 2017), *SHANK2 and SMARCA1*(Homann et al., 2016), *ANKK1*(Shirzad et al., 2016), *RBM12* (Steinberg et al., 2017), *TAAR1*(John et al., 2017), *TIMP2* (John et al., 2018)and *PTPRA*(John et al., 2018).These insightful studies suggest the importance of such variants with large/moderate effects in disease development or in increasing the risk in the respective families with disease clustering.

Considering the generally accepted polygenic/multi threshold genetic models of disease proposed to explain the disease transmission (Essen-Möller, 1977; Gershon, 2000; McGue et al., 1983; O’Rourke et al., 1982) and reported burden of rare variants in SZ patients from case-control studies (Genovese et al., 2016; Purcell et al., 2014; Richards et al., 2016), it may be hypothesized that, several uncommon variants in several genes with moderate effects may cluster in individuals in families and thus collectively contribute to disease development. This tenet may be more acceptable if we combine the reports suggesting the relatives of affected individuals showing schizotypal features and/or other endophenotypes (Breton et al., 2011; Galindo et al., 2016; Kuha et al., 2007; Lui et al., 2018; Snitz et al., 2006). Such nuclear families may manifest a pseudo Mendelian mode of disease transmission. We have recently reported a cumulative contribution of multiple protein sequence altering inherited rare variants in genes from nurodevelopmental pathways which has provided plausible genetic explanation for disease transmission and also supported the oligogenic hypothesis (John et al., 2019). Finally, considering the highly genetically heterogeneous nature of the disease (Beckmann and Franzek, 2000; Rees et al., 2015; Takahashi, 2013), such risk conferring genes may fall in a few different disease relevant pathways with their number and combinations varying across different families.

With this background, in the current study we analysed 11 multiplex SZ families by WES. This strategy facilitated identification of rare variants which may have notable functional effects and not generally captured in common variant based association studies. Multiple variants in genes which were previously implicated in SZ were observed across the eleven families supporting the polygenic/oligogenic contribution to the disease. Most of the variant/genes identified were private to each of the families reiterating the genetic heterogeneity, but they were from previously implicated pathways and gene set. Of all these, co-occurrence of several genes involved in glutamatergic signalling in affected individuals across families was most noteworthy with likely therapeutic implications.

## Methodology

### Sample recruitment

In this study 11 non consanguineous families with multiple members affected with SZ were recruited from Dr. RML Hospital New Delhi, following the diagnosis and sample recruitment procedure detailed previously(John et al., 2018, 2017, 2016; Kukshal et al., 2013). Details of the families used in the study are given in Supplementary Figure 1. Genomic DNA was extracted from peripheral blood samples from each of the participating individuals using the phenol chloroform method. All the study protocols were approved by the institutional ethical committee of Dr. RML Hospital and University of Delhi South Campus, New Delhi. Informed consent was obtained from all the participating individuals.

### Whole exome sequencing

A minimum of three members from each of the 11 families were used for WES, with preference given to affected individuals in order to obtain shared variants. Agilent SureSelect Human All Exon V5+UTR Kit was used for capture and enrichment of exomes and library preparation; the exome libraries were sequenced on Illumina HiSeq2000(101bp paired-end reads) using a commercial facility (Medgenome Labs Pvt. Ltd. Bengaluru, India, https://www.medgenome.com/).

### Sequence alignment and variant calling

Adaptors were removed from the raw sequence data in FastQ format obtained from the service provider using cutadapt(Martin, 2011). For sequence alignment and variant calling we followed the GATK(version 3.5) Best Practices workflows for germline variant discovery, as described previously(John et al., 2018, 2017). Briefly, the adaptor removed reads were aligned to the reference genome (hg19) using Burrows-Wheeler Aligner (BWA) and subsequently, PCR duplicates were removed by Picard. Using GATK realignment around insertions and deletions (indels), base quality score recalibration and variant calling were performed.

### Variant prioritisation

Kggeseq(Li et al., 2012)was used for functional annotation of variants (according to hg19 RefSeq transcripts). As per recommendations from three previous publications (Dashti and Gamieldien, 2017; MacArthur et al., 2014; Richards et al., 2015), Kggseq was also used for variant prioritisation (Li et al., 2012). As the aim of the study was to identify rare highly penetrant protein sequence altering variants we removed all the non-coding variants from the list. We removed all the common variants with minor allele frequency (MAF)>0.01 reported in public databases including 1000 genome (1000G), Exome Aggregation Consortium (ExAC r0.3.1), dbSNP141, Genome Aggregation Database (gnomAD) browser and NHLBI GO Exome Sequencing Project (ESP), along with synonymous variants. Variants shared among the affected individuals for whom exome sequencing data were available in the respective families were then assessed for segregation in the remaining available members of the respective families. Subsequently the variants which were not shared among all affected individuals in each of the families, variants from genes known to likely contribute false positive signals during variant calling (Fuentes Fajardo et al., 2012), regions with segmental duplication, which are also known to produce false positive signals and all the common variants (MAF >0.01) present in the ethnicity matched in-house data were removed. Of the remaining variants we selected start-lost, stop-gain and indels leading to frameshift or non-frameshift, canonical splice variants and all the deleterious missense variants (classified as damaging/deleterious by at least one in *silico* tool from among SIFT, Polyphen2_HDIV, Polyphen2_HVAR, LRT, MutationTaster, MutationAssessor, FATHMM, PROVEAN, MetaSVM, M-CAP and fathmm-MKL_coding and CADD scaled score =>15; or CADD scaled score>=10 and <15 but predicted to be damaging by at least two different software listed above) for further analysis. All these tools were part of Kggseq (Li et al., 2012). All these variants are henceforth referred to as “deleterious shared variants”.

### Pathway analysis and gene set enrichment analysis

In order to check the genes encompassing the “deleterious shared variants” from the 11 families, for over representation/enrichment in pathways and gene ontology categories reported to be involved in disease development, we performed pathway and gene ontology analysis using ConsensusPathDB-human (http://cpdb.molgen.mpg.de/CPDB/rlFrame) and corrected for multiple testing. To establish their enrichment in previously reported potential SZ susceptibility genes, we performed an enrichment analysis against gene sets potentially related to disease pathophysiology by selecting gene sets detailed below. Genes with *de novo* protein sequence altering variants reported in SZ (n=643)selected from denovo-db(Turner et al., 2017) (http://denovo-db.gs.washington.edu/denovo-db/index.jsp) & NPdenovo(Li et al., 2016) (http://www.wzgenomics.cn/NPdenovo/index.php); all genes reported/mapped to loci identified by GWASs of SZ (n=1683) (https://www.ebi.ac.uk/gwas/)

Since various psychiatric disorders are previously reported to have shared genetic etiology (Jensen and Girirajan, 2017; Lee et al., 2013; Martin et al., 2017), we then checked for over representation/enrichment genes involved in such disorders using the following gene sets. Genes with *de novo* protein sequence altering variants reported in, autism (n=4484); bipolar disorder (n=51) & intellectual disability (n=885) selected from denovo-db(Turner et al., 2017) & NPdenovo(Li et al., 2016); all genes reported/mapped to loci identified by GWAS of autism (n=106) and bipolar disorder (n=906)(https://www.ebi.ac.uk/gwas/home); curated list of genes reported to be involved in autism (n=990) in SFARI gene (https://www.sfari.org/resource/sfari-gene/);curated genes of bipolar disorders(n=1192) from BDgene (http://bdgene.psych.ac.cn/index.do); curated genes of psychiatric disorder (DOID_2468) (n=325) from Phenocarta (https://gemma.msl.ubc.ca/phenotypes.html), genes known to manifest abnormal behavioural/neurological phenotypes (n=3455); nervous system phenotypes (n=3528) during mouse gene knock out experiments from mouse genomics informatics(MGI). Genes (n=297) used as ASD mouse models (https://gene.sfari.org/database/animal-models/genetic-animal-models/).

In addition we also used the gene sets which are previously implicated in various psychiatric disorders including SZ and used in similar studies previously (McCarthy et al., 2014). These include loss of function intolerant genes (n=3230) from ExAC v2, 7); essential genes (n=1732)(Blake et al., 2011; Iossifov et al., 2014); FMRP interactor targets Darnell (Jennifer C. Darnell et al., 2011)(n=788) and Ascano (Ascano et al., 2012) (n=939). We used hypergeometric distribution test to calculate the significance of overlaps between “deleterious shared variants” encompassing genes and each of the gene sets mentioned above. A set of 21552 genes present in the Agilent V5 + UTR exome capture kit were used as background genes (https://www.agilent.com/en/products/genomics-agilent).

## Results

A total of 11 multimember affected SZ families were used in this study. In two families (#4 and 11), mother and her children were affected with SZ; in Family #5 father was diagnosed with bipolar disorder and mother had psychosis and their two children had SZ; and in the remaining eight families both parents didn’t show psychiatric disorders and only their children were affected with SZ and/or other psychiatric disorders (Supplementary Fig.1). A minimum of three individuals each from 11 multiplex SZ families were exome sequenced and the mean target depth of sequencing, observed across total 34 samples was 58.82X. On an average >97% of the target regions were with 10X and >89% with 20X coverage. The mean mapping quality, observed across the samples was 46.99. Using the step by step rationale for prioritization of variants detected by WES in all the selected families, as detailed in the methods section, a total of 785 deleterious shared variants in 743 genes were identified (supplementary table 1). Distribution of these variants in the respective families along with all their annotations is shown in supplementary tables 2a to 2k.

### Homozygous and compound heterozygous variants

Among the 785 rare variants, we observed only five homozygous variants in five different genes namely *PPM1K, CXCL13* and *ATAD2* in family #3, *CASC1* in family #5 and *CYP4F12* in family #9 (supplementary tables 2c, 2e and 2i).

### Variants shared across two different families

We observed one variant each in nine different genes shared across two different families. These included variants in *DCLRE1A, FRMD8, NHLRC2, PLCB3* and *SNX32* in family#5 &#7, *INPP4A, PRRT2* in families#6&#8, *PXDNL* in families#1&#11;and *ZNF730* in families#7&#8(supplementary table 3). Of note, three of the genes namely *INPP4A, PLCB3*and*PRRT2*are from the glutamatergic pathway with intronic SNP (rs12617721 p=7×10^−6^) in *INPP4A*having been reported previously in GWAS of mood-incongruent psychotic bipolar disorder (Goes et al., 2012).

### Shared gene but different variants in more than one family

We observed a total of 28 genes, but with different variants present in more than one family (supplementary table #3). Among these genes *CENPE, LAMA1* and *GRIN3B* were previously implicated in various neuro psychiatric disorders including SZ(Hornig et al., 2017; McKenna et al., 2018; Moen et al., 2017; Tarabeux et al., 2011; Xu et al., 2012). The observation of limited gene sharing across families strongly supports the commonly accepted genetic heterogeneity in SZ.

These variants were inherited from unaffected parent(s) and/or shared with unaffected members in the respective families and out of eight families where both parents were unaffected, only two families (#3 and #9 mentioned above) had homozygous variants. Considering heterozygous variants cannot give any insight based on the apparent inheritance pattern on Mendelian principles in most of the study families, we explored an alternative hypothesis.

### Functional clustering of “deleterious shared variants”

Since the monogenic disease models were inadequate to explain the disease transmission in most of the study families, we considered the alternative and commonly accepted polygenic model. We focused on the already reported genes and pathways and gene-sets in various psychiatric disorders. To evaluate the polygenic models we performed over representation/enrichment analysis using publically available pathays/GOs and gene-sets and we hypothesised that the genes from the disease relevant pathway/GO and gene-sets will be overrepresented among gene encompassing deleterious shared variants in the families.

The “deleterious shared variants” (N=785) in 743 genes (supplementary table 1) were subjected to GO and pathway analysis using ConsensusPathDB, but keeping a minimum requirement of five overlapping genes between the test set (743 genes) and reference pathways. Significant gene enrichment in extra cellular matrix related GOs; development, especially neurodevelopment related GOs; neuronal and synapse related GOs and social behaviour, axon guidance and glutamate receptor signalling pathways were some of the most striking/relevant observations, following False Discovery Rate(FDR) corrections. Of note, these findings are consistent with leading pathophysiological hypotheses of the disease (Table 1 & supplementary table 4A and 4B).

**Tabl 1:** Enriched pathways and ontologies.

**Tabl 1: Enriched pathways and ontologies**
|  |  |  | Genes in the study (family based; n=743) |  |  |
| --- | --- | --- | --- | --- | --- |
| Pathway/GO name | size | effective_size | p-value | q-value | Genes shared |
| Extracellular matrix organization | 293 | 293 | 3.36185E-05 | 0.003 | 26 |
| ECM-receptor interaction - Homo sapiens (human) | 82 | 82 | 0.0002 | 0.011 | 11 |
| Axon guidance | 357 | 356 | 0.0003 | 0.01 | 27 |
| MECP2 and Associated Rett Syndrome | 63 | 63 | 0.0005 | 0.02 | 9 |
| Wnt Signaling Pathway | 114 | 114 | 0.001 | 0.02 | 12 |
| Microtubule-based process | 776 | 776 | 6.98E-10 | 1.15E-07 | 64 |
| Cytoskeleton | 2141 | 2135 | 4.93E-09 | 6.65E-07 | 128 |
| Movement of cell or subcellular component | 2015 | 2012 | 2.24E-08 | 1.85E-06 | 120 |
| Cellular component organization | 6340 | 6325 | 4.36E-08 | 2.40E-06 | 296 |
| Microtubule associated complex | 150 | 150 | 1.13E-07 | 5.50E-06 | 21 |
| Extracellular matrix | 482 | 481 | 3.80E-07 | 1.23E-05 | 41 |
| Cytoskeletal protein binding | 916 | 914 | 1.57E-07 | 1.25E-05 | 65 |
| Cytoskeleton organization | 1318 | 1316 | 1.53E-08 | 1.40E-05 | 88 |
| Cytoskeletal part | 1636 | 1632 | 2.17E-07 | 1.62E-05 | 99 |
| Extracellular matrix component | 119 | 119 | 1.36E-06 | 3.30E-05 | 17 |
| Anatomical structure development | 5749 | 5735 | 2.02E-06 | 4.84E-05 | 264 |
| Multicellular organism development | 5261 | 5247 | 2.06E-06 | 4.84E-05 | 245 |
| Synaptic membrane | 318 | 318 | 5.30E-06 | 8.57E-05 | 29 |
| Postsynaptic membrane | 245 | 245 | 3.29E-06 | 9.82E-05 | 25 |
| Neuron part | 1538 | 1535 | 1.07E-05 | 0.0001 | 88 |
| Postsynapse | 452 | 451 | 2.09E-05 | 0.0002 | 35 |
| Neuron projection | 1228 | 1226 | 5.58E-06 | 0.0004 | 75 |
| Dorsal/ventral axis specification | 17 | 17 | 1.12E-06 | 0.0005 | 7 |
| System development | 4667 | 4655 | 6.07E-06 | 0.0005 | 219 |
| Cell development | 2039 | 2035 | 1.08E-05 | 0.0006 | 110 |
| Receptor complex | 375 | 375 | 0.0001 | 0.0008 | 29 |
| Animal organ development | 3405 | 3397 | 1.71E-05 | 0.0008 | 166 |
| Neurotransmitter receptor complex | 51 | 51 | 7.94E-05 | 0.002 | 9 |
| Axoneme part | 34 | 34 | 0.0002 | 0.003 | 7 |
| Head development | 744 | 742 | 7.77E-05 | 0.003 | 48 |
| Social behavior | 51 | 51 | 7.94E-05 | 0.003 | 9 |
| Cellular developmental process | 4298 | 4287 | 0.0004 | 0.004 | 193 |
| Nervous system development | 2304 | 2299 | 5.09E-05 | 0.005 | 118 |
| Neuron development | 1062 | 1061 | 4.31E-05 | 0.005 | 64 |
| Neurogenesis | 1541 | 1538 | 5.75E-05 | 0.005 | 85 |
| Neuron to neuron synapse | 236 | 235 | 0.0010 | 0.005 | 19 |
| Ionotropic glutamate receptor complex | 49 | 49 | 0.0003 | 0.007 | 8 |
| Asymmetric synapse | 233 | 232 | 0.0008 | 0.008 | 19 |
| Postsynaptic specialization | 230 | 229 | 0.002 | 0.008 | 18 |
| Dorsal/ventral pattern formation | 95 | 95 | 0.0002 | 0.010 | 12 |
| Axon | 540 | 540 | 0.0007 | 0.01 | 35 |
| Cell differentiation | 4103 | 4092 | 0.0004 | 0.01 | 185 |
| Tissue development | 1941 | 1938 | 0.0004 | 0.01 | 98 |
| Neuron differentiation | 1308 | 1305 | 0.0002 | 0.01 | 72 |
| Postsynaptic density | 228 | 227 | 0.002 | 0.01 | 18 |
| Cell morphogenesis involved in differentiation | 704 | 703 | 4.16E-05 | 0.01 | 47 |
| Neuron projection development | 938 | 937 | 4.90E-05 | 0.01 | 58 |
| Regulation of multicellular organismal process | 2856 | 2848 | 0.0005 | 0.01 | 135 |
| Generation of neurons | 1442 | 1439 | 7.95E-05 | 0.01 | 80 |
| Developmental growth | 584 | 584 | 0.001 | 0.01 | 36 |
| Positive regulation of developmental process | 1265 | 1263 | 0.0004 | 0.02 | 69 |
| Brain development | 703 | 701 | 0.0006 | 0.02 | 43 |
| Developmental maturation | 256 | 256 | 0.003 | 0.02 | 19 |
| Dendrite development | 217 | 216 | 0.0009 | 0.02 | 18 |
| Positive regulation of nervous system development | 493 | 491 | 0.0002 | 0.02 | 34 |
| Animal organ morphogenesis | 996 | 995 | 0.001 | 0.03 | 55 |
| Embryo development | 956 | 956 | 0.001 | 0.03 | 53 |
| Neural precursor cell proliferation | 136 | 136 | 0.004 | 0.03 | 12 |
| Regulation of developmental process | 2456 | 2450 | 0.002 | 0.03 | 115 |
| Regulation of localization | 2615 | 2610 | 0.003 | 0.05 | 120 |
| Developmental process involved in reproduction | 669 | 669 | 0.007 | 0.05 | 37 |
| Neural tube development | 158 | 158 | 0.002 | 0.05 | 14 |
| Receptor clustering | 50 | 50 | 0.002 | 0.05 | 7 |
| Regulation of multicellular organismal development | 1897 | 1891 | 0.002 | 0.05 | 92 |

Details of all other enrichment analysis based on seemingly relevant and previously used gene-sets are presented in supplementary table 5 and are briefly presented below. Although no enrichment of genes from SZ, autism and bipolar disorder GWASs was documented, an overlap of 46, 5 and 25 genes respectively, from these three GWASs was noted. Among these, *ANK3* and *PPM1M* were reported in both SZ and bipolar disorder GWASs. On the contrary, significant overrepresentation of genes encompassing *de novo* mutations reported in SZ (p= 2.92E-05); autism (p= 2.35E-18) and intellectual disability (p= 7.87E-04) were observed. In addition, enrichment of genes reported in related disease databases, like bipolar disorder (p=0.001, from BDgene database); autism (p=5.014E-07, from SFARI Gene database); psychotic disorder (p=6.65E-05 from Phenocarta) were observed. A few other notable findings included FMRP targets including Darnell and Ascano (p=6.258E-06 and p=1.617E-06 respectively), and Exac_LOF_Intolerant genes (p=1.315E-04).

On analysis with the mouse genomics informatics (MGI) we observed a total of 163 genes among the 742 which were known to manifest “behavioural/neurological phenotype” during knock out studies and which were overrepresented in our studies (P=1.09E-05). In addition 19 genes were used previously as animal models in autism studies. Manual literature search also identified a total of 52 genes known to show schizophrenia relevant phenotypes during inactivation or over expression in various rodent models (Supplementary text file). In this way a total of 175 genes showed various psychiatry relevant phenotypes during rodent knockout/over expression studies. All these results reiterate that the “deleterious shared variants” encompassing genes are enriched with SZ and other psychiatry relevant candidate genes and genes from the disease relevant pathways and GOs/Gene-set.

### Genetic heterogeneity and polygenic nature

Since the variants/ (9/785;∼1%)/genes (28 /734; ∼4%) that were shared in more than one family was ∼5%, it suggests that variants/genes are mostly private to each family and supports genetic heterogeneity. Enrichment of genes involved in various SZ relevant pathways and genes from various gene-sets previously implicated in SZ and psychiatric disorders; a large number of genes which showed behavioural abnormalities in various rodent models; and differential expression in various psychiatric disorders highlight the polygenic nature of the disease.

### Identification of most promising risk conferring variants in the study families

Findings from the enrichment analysis detailed above, motivated us to review the individual family data. Based on leading pathophysiological hypothesis and current knowledge about disease biology, we undertook to check whether disease relevant genes) with the rare variants are notably more clustered in the affected individuals in each of the families.

As an exploratory analysis we prioritised the “deleterious shared variants” in each family based on their previous genetic reports, reported pathophysiological hypothesis and evidence from animal models. Through this approach we identified a total of 122 variants from 117 genes (supplementary tables2 & 6) as the most promising risk conferring genes in the 11 families, with an average of ∼11 variants per family. 82 of these genes were previously reported in SZ/bipolar disorder/autism GWASs or *de novo* mutation studies of SZ/bipolar disorder/autism/intellectual disability and 42 genes have been reported in mouse knockout/over expression studies wherein psychiatry relevant behavioural abnormalities were documented. These included *CALHM1, ANK2, ANK3, CHRM4, DISC1, DLG1, DRD3, EGF, GRIN3A, GRM2, GRM7, HTR3A, KALRN, KCNT1, LRP4, MAGI2, PRODH, SCN2A, SETDB1, SHANK2, SHANK3, TBX1, TH, WNT2, PLCB3, INPP4A, PRRT2, CENPE, LAMA1* and *GRIN3B.* Most promising risk conferring genes thus identified have been presented family wise in Supplementary Figs 1a….1k; Supplementary Table 2. Unaffected members also shared these variants, however the total variants present in the unaffected members were half the number compared to the affected individuals. Unaffected parents transmitted the rare variants almost equally to the affected progeny. Known functions and all other available supporting evidence including from animal studies for likely involvement of genes in disease biology are presented in Supplementary text file. This is not to assign the complete genetic basis of the disease in each individual or to claim that these genes/variants are the only contributors to disease etiology, but rather to check the distribution of the most promising rare variant encompassing candidate genes across the affected and unaffected individuals in each of the families, keeping in mind their documented probable functional relevance and pathophysiological hypothesis. This was also done to check if such an approach could help interpret the complex mode of inheritance with familial clustering of psychiatric disorders which characterised the study families. Original findings of negligible contribution of homozygous/compound heterozygous variants in SZ families were parents were unaffected, and heterozygous variants in large number of disease relevant genes following this analysis (Supplementary Fig1) provides genetic evidence for cumulative contribution of multiple rare heterozygous variants as well as a likely threshold effect for disease manifestation.

## Discussion

Uncovering the genetic underpinnings of neuropsychiatric disorders have witnessed distinct eras starting with limited hypothesis testing/candidate gene association testing in small/moderate sample sizes, through hypothesis free genome wide analysis of common variants across large cohorts including trans-ethnic groups, to the contemporary rare variant paradigm. Rare variant findings have provided significant insights into several disease related pathways but the variants themselves were mostly *de novo* in origin with limited scope to explain genetic liability across generations. Conversely, analysis of multiplex families with the same rare variant discovery tool of whole exome/genome sequencing is expected to be more powerful to detect rare inherited rare protein sequence disrupting variants which maybe potentially more useful in genetic risk prediction.

Results from our efforts in this direction by analysing 11 small sized families have been encouraging and have underscored the contribution of inherited risk variants to disease etiology. The notable observations included: a large number of inherited rare (MAF <0.01) protein sequence altering and deleterious variants in a large number of disease relevant genes (including well studied SZ candidate genes such as *CALHM1, ANK2, ANK3, CHRM4, DISC1, DLG1, DRD3, EGF, GRIN3A, GRM2, GRM7, HTR3A, KALRN, KCNT1, LRP4, MAGI2, PRODH, SCN2A, SETDB1, SHANK2, SHANK3, TBX1, TH* and *WNT2*) across the 11 multiplex SZ families; variants in 39 genes (including genes such as *INPP4A, CENPE, LAMA1* and *GRIN3B*) present in two families; enrichment of genes from disease relevant pathways/ontologies including neurodevelopmental, synaptic functions, social behaviour, axon guidance, glutamatergic pathway etc. (Table 1; Supplementary Table 4); and notable clustering of genes with previously reported *de novo* mutations in SZ as well as other psychiatric disorders (Supplementary Table 5). In addition, most of the variants were heterozygous (Suppl_Tables_2A-to-2K) and this is not unexpected considering their rarity and non-consanguineous nature of the study families. Furthermore, occurrence of rare variants in functionally prioritised genes (Supplementary Table 6) was noteworthy. Most of the variants were directly/indirectly connected with glutamatergic pathways (Supplementary text file). Of note, at least five genes each in all the families were reported previously to manifest behavioural and nervous system abnormalities in mouse knock out studies. A brief description of functionally prioritised genes is provided in Supplementary text file.

Highlights which emerge from these family based findings are i) the strong genetic evidence that it has provided for the cumulative contribution of protein sequence altering rare variants to disease development; ii) identification that unaffected members possessed only about half the total number of the inherited rare variants documented in the affected members in the respective families (Supplementary Fig1), suggesting a threshold effect; and iii)a likely explanation for the complex mode of inheritance seen in the study families. Furthermore, the limited extent of overlapping genes across the families reiterates the genetic heterogeneity reported in SZ. Almost all of the prioritised genes in each of the families were relevant to psychiatric disorders in general as evident in published studies (Supplementary information).

The most significant enrichment of pathways and GOs observed in the analysis is related to microtubule/extracellular matrix/cytoskeleton related GOs and pathways(Table 1 & Supplementary Table 4).Cytoskeleton is the basic structural framework of the cell and comprised of microtubules, actin filaments, and intermediate filaments. In general it is involved in maintenance of cell shape and internal organization through linkages to itself, the membrane, and internal organelles. Cytoskeleton in brain is involved in development, maintenance and regeneration of neurons (see review (Yan et al., 2016)). It is also involved in spine development, receptor anchoring and trafficking, trans-synaptic adhesion and structural plasticity during long term potentiation and depression (Forsyth and Lewis, 2017). The authors proposed that synaptic actin dysfunction may be the convergent mechanism of various psychiatric disorders. NMDA receptor signalling influences the reorganisation of cytoskeleton (Forsyth and Lewis, 2017). Microtubule dysfunction has been previously reported in SZ and other psychiatric disorders (Gardiner, 2017; Marchisella et al., 2016). An over representation of *de novo* mutation encompassing genes from activity-regulated cytoskeleton-associated protein complex in SZ has been previously reported (Fromer et al., 2014). On the other hand, abnormalities in extracellular matrix (ECM) are also reported in SZ and other psychiatric disorders. ECMs are involved in synaptic functions, neuronal connectivity and migration, and GABAergic, glutamatergic and dopaminergic neurotransmission(Berretta, 2012; Matuszko et al., 2017). Another ontological process where enrichment was observed is developmental processes especially brain development, supporting the widely accepted neurodevelopmental models of SZ (Owen et al., 2011; Owen and O’Donovan, 2017). Enrichment of genes from other related pathways/GOs include ionotropic glutamate receptor complex, glutamate receptor related pathways/GOs and N-methyl-D-aspartate glutamate receptor (NMDAR).

A notable over representation was seen in FMRP. This RNA binding protein is coded by *FMRI* (Xq27.3) which is linked to fragile X syndrome. FMRP is important for translation of hundreds of neuronal mRNAs and affects synaptic functions (Ascano et al., 2012; Jennifer C. Darnell et al., 2011). Taken together, over representation of genes reported from the *de novo* mutation studies in autism, ID and other curated list of genes involved in other psychiatric disorders (taken from various databases) (Supplementary Table 5), lend support to a likely shared etiology across these brain disorders(Gandal et al., 2018; Lee et al., 2013; O’Donovan and Owen, 2016; Smoller et al., 2013; Zhao and Nyholt, 2017).

The above discussed study findings are indeed informative and support a polygenic nature of SZ, but have a few limitations. i) In view of the large number of inherited variants, only those predicted to be damaging by *in silico* tools have been considered which may lead to a few functionally relevant variants being missed; ii) *de novo* variants in the families have not been assessed; iii) focus has been only on rare SNPs/indels thus ignoring CNVs and common variants; iv) considering only those variants shared among all SZ affected in the respective families may have led to omission of private variants which may be otherwise potential contributors; similarly non coding variants have also not been considered; and finally, v) only small sized families have been analysed which may imply suboptimal power of the study. However, considering the overarching findings of inherited rare variants in genes of functional relevance to SZ segregating among the affected individuals in this study, efforts to investigate larger multi ethnic family based and case-control cohorts on one hand and functional validation of the variants using *in vitro* or cellular/animal models on the other would be insightful. Further, weighted polygenic scoring considering genes of biological relevance and nature and genomic location of the variants with respect to brain function would be desirable for complex diseases such as SZ.

## Acknowledgements

Junior and Senior Research Fellowship (09/045(1166)/2012-EMR-I) to Jibin John from Council for Scientific and Industrial Research (CSIR), New Delhi; and DSK-PDF (BL/13-14/0404) to Dr. Prachi Kukshal from UGC, New Delhi are gratefully acknowledged. We are thankful for, the study sample collection by trained and dedicated staff at Dr RML hospital, DNA isolation by Mrs. Anjali Dabral at the University of Delhi South Campus; and computational facility provided by Central Instrumentation Facility, University of Delhi South Campus. We gratefully acknowledge infrastructure support provided by the UGC, New Delhi, through Special Assistance Programme and Department of Science and Technology, New Delhi, through FIST and DU-DST PURSE programmes to the Department of Genetics, UDSC.

