## Supplemental Text for "Oligogenic rare variant contributions in schizophrenia and their convergence with genes harbouring *de novo* mutations in schizophrenia, autism and intellectual disability: Evidence from multiplex families"

**Supplemenary information: Functions of prioritised genes**

### Family #1

**Family #2**

1. **GABRR3:** This gene encodes a subunit of the GABA(C) receptor. Gamma amino butyric acid (GABA) is one of the neurotransmitters in the central nervous system and it regulates synaptic neurotransmission in the neurons. Alteration in the GABA neurotransmitter system is observed in SZ (Wassef et al., 2003)
2. **CASR**: involved in synaptic plasticity, neurotransmission, neuronal growth (Ruat and Traiffort, 2013) and promotes resting spontaneous glutamate release(Ruat and Traiffort, 2013).
3. **DOCK3:** regulates BDNF-TrkB signalling and, neurite and axonal outgrowth (Namekata et al., 2012, 2010). It interacts with NMDA receptors containing NR2D subunit and reduces the surface expression of NR2D (Bai et al., 2013)
4. **PDE8B:** encodes cyclic nucleotide phosphodiesterase (PDE). It is involved in the hydrolysis of the second messenger cAMP and is involved in various cognitive functions (Tsai et al., 2012)
5. **MAGI2:** involved in the recruitment of AMPA and NMDA-type glutamate receptors (Deng et al., 2006). Structural variations reported in patients with SZ and bipolar disorder (Karlsson et al., 2012; Walsh et al., 2008). Common variants in this gene reportedly increased risk of cognitive impairment in patients with SZ (Koide et al., 2012).
6. **GLI3:** involved in development of the cerebral cortex, hypothalamus, corpus callosum (Amaniti et al., 2013; Haddad-TÃ³volli et al., 2015; Wilson et al., 2012), and maintenance and fate specification of cortical progenitors (Wang et al., 2011)
7. **VPS13B: Has an** important role in the development of the central nervous system
8. **CALHM1:** involved in the regulation of $Ca^{2+}$-dependent MEK, ERK, RSK and MSK signalling in cerebral neurons (Dreses-Werringloer et al., 2013). Involved in memory, long-term potentiation and NMDA and AMPA receptors phosphorylation **and**

**knockout mice showed a severe impairment in memory flexibility**(Vingtdeux et al., 2016)

9. **INA:** is involved in intracellular transport to axons and dendrites and morphogenesis of neurons.
10. **CACNG7:** encodes a type II transmembrane AMPA receptor regulatory protein (TARP), involved in synaptic expression of cerebellar AMPA receptors and function (Kato et al., 2007; Studniarczyk et al., 2013; Yamazaki et al., 2010)
11. **PLAUR**: encodes the receptor for urokinase plasminogen activator and the gene is involved in the development of GABAergic interneurons and **knockout mice have shown various behavioural abnormalities** (Eagleson et al., 2011, 2010)

**Family #3**

8. **NTN1:** is a member of laminin-related secreted proteins. Netrin-1 is implicated in the organization, plasticity and function of mesocorticolimbic DA neurons in rodents(Flores, 2011).

**Family #4**

1. **LRRC7:** It is known to influence the function of mGluRs and CaMKII at synapses and involved in the localisation of mGluR5 and DISC1 in the PSD fraction. **Knockout mice**

**showed various relevant endophenotypes of schizophrenia and autism spectrum disorders**(Carlisle et al., 2011)**.**

**Family #5**

1. **KDM4A:** is involved in BDNF expression, neuronal differentiation and survival (Milstein et al., 2007).
2. **NOTCH2**: is a member of the Notch family and is involved in various developmental processes. In hippocampal neurons, NOTCH2 is involved in the regulation of Vesicular glutamate transporter 1 (VGLUT1) protein levels (Wojcik et al., 2004)
3. **DLG1:** is a multi-domain scaffolding protein involved in various developmental processes. In the brain the protein is involved in trafficking and localisation of AMPA and NMDA-type glutamate receptor, Kv1.4, Kv4.2, and Kir2.2 potassium channels and astrocytic glutamate transporter EAAT2b (Cai et al., 2006; Fourie et al., 2014; Hong et

al., 2015; Underhill et al., 2015; Walch, 2013). **Knockout mice showed reduced sociability** (Coba et al., 2018).

**Family #6**

**impairment in rearing activity as well as in fear-conditioned memory and significantly reduced levels of serotonin levels in the plasma and brain** (Tuoc et al., 2009).

**Family #7**

1. **DISC1:** DISC1 is proposed to be involved in the release of glutamate (Maher and LoTurco, 2012) and known to impact the NMDAR expression and function (Yan et al., 2014). ***DISC1* knockout mouse models showed various phenotypes relevant to schizophrenia and other psychiatric disorders** (Jaaro-Peled, 2009; Johnstone et al., 2011)
2. **AGRN:** plays an important role in neuronal responses to excitatory neurotransmitters both *in vitro* and *in vivo* (Hilgenberg et al., 2002). Agrin also has an important role in synaptic development, synaptic activity and plasticity (Daniels, 2012)
3. **PTPRN:** member of the protein tyrosine phosphatase (PTP) family, required for normal accumulation of the neurotransmitters norepinephrine, dopamine and serotonin in the brain. **Knockout mice showed various behavioural and cognitive abnormalities** (Cai and Notkins, 2016; Carmona et al., 2014; Nishimura et al., 2010, 2009)
4. **SLC25A12:** encodes a calcium binding mitochondrial protein and it is involved in exchange of aspartate for glutamate across the inner mitochondrial membrane. The

gene has been previously implicated in both SZ and autism (Aoki and Cortese, 2016; Hong et al., 2007). **Knockout mice showed altered dopamine levels in the brain, hyper-reactivity and anxiety-like behaviour** (Llorente-Folch et al., 2013)

5. **WNT2:** involved in the development of dopaminergic progenitors and dopaminergic neurons. **Knockdown of the gene in mouse brains cause depression-like behaviours, impaired Wnt/β-catenin signalling and neurogenesis** (Sousa et al., 2010; W. J. Zhou et al., 2016)
6. **PTPRD:** is a member of the protein tyrosine phosphatase (PTP) family and the gene is involved in hippocampal long-term potentiation and learning processes (Uetani, 2000). Recent GWAS showed that PTPRD is associated with Antipsychotic-induced weight gain (Yu et al., 2016). **PTPRD deficient mice showed impairment in cognition and enhanced long-term potentiation (LTP)** (Bruining et al., 2015). Structural variation is reported in autism spectrum disorder, dipolar disorder and ADHD (Bruining et al., 2015)
7. **PLCB3:** encodes a member of the phosphoinositide phospholipase C beta enzyme family. Through the interaction with SHANK2, PLCB3 regulates mGluR-mediated calcium signalling in the brain (Hwang et al., 2005)
8. **PCDH9**: is a member of the protocadherin family and the gene is involved in various neurodevelopmental processes (Asahina et al., 2012). **Knockout mice showed specific long-term social and object recognition deficits** (Bruining et al., 2015)
9. **DYNC1H1:** is a microtubule activated ATPase and acts as a molecular motor. This gene is involved in the modulation of dopamine signalling in striatum (Braunstein et al., 2010) and various neuro developmental processes (Ori-McKenney and Vallee, 2011).
10. **TRPV3:** is a member of nonselective cation channels and involved in temperature sensation and vasoregulation. Recently its role in modulation of mesolimbic-dopamine signalling has been reported (Singh et al., 2016).
11. **PNPO:** encodes pyridoxamine 5'-phosphate oxidase, a rate-limiting enzyme in pyridoxal 5'-phosphate (vitamin B(6)) synthesis. Vitamin B6 acts as a co-factor in the synthesis of neurotransmitters such as catecholamine and in homocysteine metabolism.
12. **BSG:** is a plasma membrane protein. It is an obligatory subunit of native plasma membrane $Ca^{2+}$-ATPases,is involved in the extrusion Ca2+ ions from the cytosol and is a regulator of $Ca^{2+}$ signalling. So it is involved in the regulation of intracellular Ca2+ concentration(Kina et al., 2007).**Knockout mice showed abnormalities in sensory and memory functions** (Naruhashi et al., 1997)

**Family #8**

1. **EIF2AK3: Through** phosphorylation, it inactivates the alpha subunit of eukaryotic translation-initiation factor 2 and thus causes repression of translational initiation and global protein synthesis. **Brain-specific disruption of the gene in mice leads to various behavioural abnormalities including cognitive and information processing deficits, and reduced prepulse inhibition** (Trinh et al., 2012)
2. **INPP4A:** encodes an enzyme which catalyses dephosphorylation of phosphatidylinositol 3,4-bisphosphate, inositol 1,3,4-trisphosphate and inositol 3,4-bisphosphate. It is found in postsynaptic density and involved in the regulation of NMDAR-mediated excitatory postsynaptic current and localization of NMDAR in the synapse (Sasaki et al., 2010).
3. **PTPN4:** is a member of the protein tyrosine phosphatase (PTP) family and in the brain

it interacts with glutamate receptor delta 2 and epsilon subunits(Hironaka et al., 2000). The gene is also involved in phosphorylation and dephosphorylation of glutamate receptors (GluD2, GluA2) and thus long-term depression (Kohda et al., 2013). It is also involved in learning and cerebellar synaptic plasticity (Kina et al., 2007).**Knockout mice had shown impaired motor learning and cerebellar long-term depression** (Kina et al., 2007).

**Family #9**

1. **SCN2A:** is a voltage-gated sodium channel involved in the generation and propagation of action potentials in excitable cells. This gene is already implicated in various

neuropsychiatric disorders including SZ (Carroll et al., 2016). ***Scn2a*$^{KO/+}$ mice showed a range of behavioural and neurological phenotype observed in various rodent models of schizophrenia and autism spectrum disorder** (Trinh et al., 2012).

.

**Family #10**

1. **PLXNA2:** is a member of the plexin-A family of semaphorin co-receptors and has a role in axon guidance, invasive growth and cell migration. ***Plxna2***$^{-/-}$ **mice showed malformed dentate gyrus and SZ like behaviours** (Zhao et al., 2018)**.**

2. **GRM7:** is a member of the metabotropic class of glutamate receptors.

3. **GRM2:**is a member of the metabotropic class of glutamate receptors. **Through positive allosteric modulation of mGluR2 in translational models of cognitive symptoms associated with schizophrenia, showed improvement in memory and attention deficits** (Yu et al., 2013)
4. **PLD1:** encodes phosphatidylcholine-specific phospholipase and catalyses the hydrolysis of phosphatidylcholine to phosphatidic acid and choline. In the brain, it is involved in BDNF induced signalling (Ammar et al., 2015), which in turn is involved in glutamatergic and GABAergic synapse formation, maturation and plasticity (Gottmann et al., 2009)**. Knockout mice displayed impaired brain development and reduced cognitive function** (Zoubovsky et al., 2015)**.**
5. **PCM1:** is a DISC1 interacting protein, implicated in centrosome assembly and function, implicated in SZ (Moens et al., 2010). **Haploinsufficient for *PCM1* (*Pcm1*$^{+/-}$) mice showed reduced whole brain volume and impairment in social interaction** (Zoubovsky et al., 2015)**.**
6. **NRP1:** is a semaphorin receptor, involved in neuronal migration and patterning (Fantin et al., 2009). During the construction of GABAergic circuitry, NRP1 is involved in axon guidance and subcellular target recognition (Telley et al., 2016).
7. **NEURL1:** Through the modulation of CPEB3 levels and CPEB3-dependent synthesis of GluR1 and GluR2, NEURL1 is involved in hippocampal plasticity and hippocampal-dependent memory (Yu et al., 2013). In addition, it also affects the growth of new dendritic spines and increases the expression of GluA1 and GluA2 subunits of AMPA receptors. **Inhibition of the gene impairs hippocampal-dependent memory and synaptic plasticity, and reduces GluA1 and GluA2 levels** (Pavlopoulos et al., 2011)**.**
8. **TH:** is a rate limiting enzyme in the biosynthesis of catecholamine, like dopamine and noradrenaline.
9. **IQSEC3:** is a member of the brefeldin A-resistant Arf-GEF/IQSEC family. Through interaction with gephyrin IQSEC3, promotes the formation of inhibitory synapses (Fukaya et al., 2011; Um et al., 2016).
10. **THBS1**: is a glycoprotein and is involved in cell-to-matrix and cell-to-cell interactions and in the regulation of excitatory synaptogenesis through the gabapentin receptor $\alpha 2\delta$-1(Crawford et al., 2012; Eroglu et al., 2009) and neuroligin 1 (NRG1)(Xu et al., 2010).
11. **LAMA1:** encodes one of the alpha 1 subunits of laminin and is involved in cell adhesion, migration, neurite outgrowth and angiogenesis. It has an important role in cerebellum development. **Knockout mice showed behavioural abnormalities and impaired formation of the cerebellum** (Heng et al., 2011; Ichikawa-Tomikawa et al., 2012)
12. **EFNA2:** in the developing neocortex, it is involved in interneuron migration and

excitatory neuron differentiation (Homman-Ludiye et al., 2017). It is also involved in the modulation of experience-dependent, NMDA receptor-mediated, synaptic pruning during maturation of the mouse cortex (Yu et al., 2013).

13. **PTPRT:** is a member of the protein tyrosine phosphatase (PTP) family. Through interaction with cell adhesion molecules and Fyn protein tyrosine kinase, PTPRT regulates synapse formation in CNS (Lim et al., 2009). (Please rewrite the next two sentences. They don't make sense as is.)Through interactions between the Syntaxin-binding protein, the Syntaxin protein known to regulate neurotransmitter release and the protein also known to be involved in various neurodevelopmental process by dephosphorylating synaptic molecules and its interaction with neuronal adhesion molecules (Lee, 2015).
14. **SHANK3:** is a multidomain scaffold postsynaptic density protein of excitatory synapses. It is involved in NMDA receptor signalling (Duffney et al., 2013), regulation of Metabotropic Glutamate Receptor 5 expression (Duffney et al., 2013) and AMPA Receptor Recycling and Synaptic LTP (Raynaud et al., 2013). **SHANK3 knockout mice showed autistic-like behaviours, striatal dysfunction, spatial memory deficit and abnormal social interaction** (Jaramillo et al., 2016; Lee et al., 2015; Peça et al., 2011)
15. **PLXNA3:** is a class 3 semaphorin receptor involved in the development of hippocampal axonal projections (Pavlopoulos et al., 2011)

**Family #11**

1. **ANK2:** is a member of the Ankyrin protein family and connects integral membrane proteins to underlying spectrin-actin cytoskeleton. The gene has an important role in synaptic stability (Koch et al., 2008).
2. **KMT2E:** involved in promoter related histone acetylation and H3K4 tri- and di-methylation
3. **FGF23:** is a member of the fibroblast growth factor family of proteins. In the hippocampal neurons, it induces phospholipase Cγ activity, inhibits neuronal ramification and enhances the synaptic density. It may also be involved in memory formation (Hensel et al., 2016).
4. **GRIN3B**: is a subunit of a glutamate N-methyl-D-aspartate (NMDA) receptor**. Knockout mice showed impairment in motor learning or coordination, changes in home cage activity, social interaction, and increased anxiety-like behaviour** (Kina et al., 2007)
5. **ARR3:** encodes a non-visual arrestin, which binds to agonist-activated, phosphorylated G protein-coupled receptors. This dissociates receptors from the heterotrimeric G protein and thus terminates the signalling process. **ARR3** is involved in D2 dopamine receptor internalisation (Skinbjerg et al., 2009).

Ito, H., Shiwaku, H., Yoshida, C., Homma, H., Luo, H., Chen, X., Fujita, K., Musante, L., Fischer, U., Frints, S.G.M., Romano, C., Ikeuchi, Y., Shimamura, T., Imoto, S., Miyano, S., Muramatsu, S.I., Kawauchi, T., Hoshino, M., Sudol, M., Arumughan, A., Wanker,

E.E., Rich, T., Schwartz, C., Matsuzaki, F., Bonni, A., Kalscheuer, V.M., Okazawa, H., 2015. In utero gene therapy rescues microcephaly caused by Pqbp1-hypofunction in neural stem progenitor cells. Mol. Psychiatry 20, 459–471. https://doi.org/10.1038/mp.2014.69

Ito, H., Yoshimura, N., Kurosawa, M., Ishii, S., Nukina, N., Okazawa, H., 2009. Knock-down of PQBP1 impairs anxiety-related cognition in mouse. Hum. Mol. Genet. 18, 4239–4254. https://doi.org/10.1093/hmg/ddp378

Jaaro-Peled, H., 2009. Gene models of schizophrenia: DISC1 mouse models. Prog. Brain Res. 179, 75–86. https://doi.org/10.1016/S0079-6123(09)17909-8

Jacquet, H., Raux, G., Thibaut, F., Hecketsweiler, B., Houy, E., Demilly, C., Haouzir, S., Allio, G., Fouldrin, G., Drouin, V., Bou, J., Petit, M., Campion, D., Frébourg, T., 2002. PRODH mutations and hyperprolinemia in a subset of schizophrenic patients, Human Molecular Genetics. https://doi.org/10.1093/HMG/11.19.2243

Jahn, R., Fasshauer, D., 2012. Molecular machines governing exocytosis of synaptic vesicles. Nature. https://doi.org/10.1038/nature11320

Janisch, K.M., Vock, V.M., Fleming, M.S., Shrestha, A., Grimsley-Myers, C.M., Rasoul, B.A., Neale, S.A., Cupp, T.D., Kinchen, J.M., Liem, K.F., Dwyer, N.D., 2013. The vertebrate-specific Kinesin-6, Kif20b, is required for normal cytokinesis of polarized cortical stem cells and cerebral cortex size. Development 140, 4672–4682. https://doi.org/10.1242/dev.093286

Jaramillo, T.C., Speed, H.E., Xuan, Z., Reimers, J.M., Liu, S., Powell, C.M., 2016. Altered Striatal Synaptic Function and Abnormal Behaviour in Shank3 Exon4-9 Deletion Mouse Model of Autism. Autism Res. 9, 350–375. https://doi.org/10.1002/aur.1529

Jensen, L.R., Chen, W., Moser, B., Lipkowitz, B., Schroeder, C., Musante, L., Tzschach, A., Kalscheuer, V.M., Meloni, I., Raynaud, M., van Esch, H., Chelly, J., de Brouwer, A.P.M., Hackett, A., van der Haar, S., Henn, W., Gecz, J., Riess, O., Bonin, M., Reinhardt, R., Ropers, H.-H., Kuss, A.W., 2011. Hybridisation-based resequencing of 17 X-linked intellectual disability genes in 135 patients reveals novel mutations in ATRX, SLC6A8 and PQBP1. Eur. J. Hum. Genet. 19, 717–720. https://doi.org/10.1038/ejhg.2010.244

Ji, X., Takahashi, N., Saito, S., Ishihara, R., Maeno, N., Inada, T., Ozaki, N., 2008. Relationship between three serotonin receptor subtypes (HTR3A, HTR2A and HTR4) and treatment-resistant schizophrenia in the Japanese population. Neurosci. Lett. 435, 95–98. https://doi.org/10.1016/j.neulet.2008.01.083

Jia, Z., Guo, Y., Tang, Y., Xu, Q., Li, B., Wu, Q., 2014. Regulation of the Protocadherin Celsr3 Gene and Its Role in Globus Pallidus Development and Connectivity. Mol. Cell. Biol. 34, 3895–3910. https://doi.org/10.1128/MCB.00760-14

Jiang, Y., Loh, Y.H.E., Rajarajan, P., Hirayama, T., Liao, W., Kassim, B.S., Javidfar, B., Hartley, B.J., Kleofas, L., Park, R.B., Labonte, B., Ho, S.M., Chandrasekaran, S., Do, C., Ramirez, B.R., Peter, C.J., Julia, T.C.W., Safaie, B.M., Morishita, H., Roussos, P., Nestler, E.J., Schaefer, A., Tycko, B., Brennand, K.J., Yagi, T., Shen, L., Akbarian, S., 2017. The methyltransferase SETDB1 regulates a large neuron-specific topological chromatin domain. Nat. Genet. 49, 1239–1250. https://doi.org/10.1038/ng.3906

Jo, K., Derin, R., Li, M., Bredt, D.S., 1999. Characterization of MALS/Velis-1, -2, and -3: a family of mammalian LIN-7 homologs enriched at brain synapses in association with the postsynaptic density-95/NMDA receptor postsynaptic complex. J. Neurosci. 19, 4189–4199. https://doi.org/1999/05/26 00:01

Johnson, S., Stockmeier, C.A., Meyer, J.H., Austin, M.C., Albert, P.R., Wang, J., May, W.L., Rajkowska, G., Overholser, J.C., Jurjus, G., Dieter, L., Johnson, C., Sittman, D.B., Ou, X.M., 2011. The reduction of R1, a novel repressor protein for monoamine oxidase a, in major depressive disorder. Neuropsychopharmacology 36, 2139–2148. https://doi.org/10.1038/npp.2011.105

Johnstone, M., Thomson, P.A., Hall, J., McIntosh, A.M., Lawrie, S.M., Porteous, D.J., 2011. DISC1 in schizophrenia: Genetic mouse models and human genomic imaging. Schizophr. Bull. 37, 14–20. https://doi.org/10.1093/schbul/sbq135

Kaeser-Woo, Y.J., Younts, T.J., Yang, X., Zhou, P., Wu, D., Castillo, P.E., Sudhof, T.C., 2013. Synaptotagmin-12 Phosphorylation by cAMP-Dependent Protein Kinase Is Essential for Hippocampal Mossy Fiber LTP. J. Neurosci. 33, 9769–9780. https://doi.org/10.1523/JNEUROSCI.5814-12.2013

Karakatsani, A., Marichal, N.N., Urban, S., Kalamakis, G., Ghanem, A., Schick, A., Zhang, Y., Conzelmann, K.-K., Rüegg, M.A., Berninger, B., de Almodovar, C.R., Gascón, S., Kröger, S., Ruiz de Almodovar, C., Gascón, S., Kröger, S., 2017. Neuronal LRP4 regulates synapse formation in the developing CNS. Development 144, dev.150110. https://doi.org/10.1242/dev.150110

Karlsson, R., Graae, L., Lekman, M., Wang, D., Favis, R., Axelsson, T., Galter, D., Carmine Belin, A., Paddock, S., 2012. MAGI1 copy number variation in bipolar affective disorder and schizophrenia. Biol. Psychiatry 71, 922–930. https://doi.org/10.1016/j.biopsych.2012.01.020

Karpyak, V.M., Geske, J.R., Colby, C.L., Mrazek, D.A., Biernacka, J.M., 2012. Genetic variability in the NMDA-dependent AMPA trafficking cascade is associated with alcohol dependence. Addict. Biol. 17, 798–806. https://doi.org/10.1111/j.1369-1600.2011.00338.x

Kato, A.S., Zhou, W., Milstein, A.D., Knierman, M.D., Siuda, E.R., Dotzlaf, J.E., Yu, H., Hale, J.E., Nisenbaum, E.S., Nicoll, R.A., Bredt, D.S., 2007. New Transmembrane AMPA Receptor Regulatory Protein Isoform, -7, Differentially Regulates AMPA Receptors. J. Neurosci. 27, 4969–4977. https://doi.org/10.1523/JNEUROSCI.5561-06.2007

Kawauchi, T., Chihama, K., Nabeshima, Y.I., Hoshino, M., 2003. The in vivo roles of STEF/Tiam1, Rac1 and JNK in cortical neuronal migration 22, 4190–4201. https://doi.org/10.1093/emboj/cdg413

Kele, J., Andersson, E.R., Villaescusa, J.C., Cajanek, L., Parish, C.L., Bonilla, S., Toledo, E.M., Bryja, V., Rubin, J.S., Shimono, A., Arenas, E., 2012. SFRP1 and SFRP2 dose-dependently regulate midbrain dopamine neuron development in vivo and in embryonic stem cells. Stem Cells 30, 865–875. https://doi.org/10.1002/stem.1049

Kim, E.J., Ables, J.L., Dickel, L.K., Eisch, A.J., Johnson, J.E., 2011. Ascl1 (Mash1) defines cells with long-term neurogenic potential in subgranular and subventricular zones in adult mouse brain. PLoS One 6, e18472. https://doi.org/10.1371/journal.pone.0018472

Kim, I.H., Racz, B., Wang, H., Burianek, L., Weinberg, R., Yasuda, R., Wetsel, W.C., Soderling, S.H., 2013. Disruption of Arp2/3 Results in Asymmetric Structural Plasticity of Dendritic Spines and Progressive Synaptic and Behavioral Abnormalities. J. Neurosci. 33, 6081–6092. https://doi.org/10.1523/JNEUROSCI.0035-13.2013

Kina, S.I., Tezuka, T., Kusakawa, S., Kishimoto, Y., Kakizawa, S., Hashimoto, K., Ohsugi, M., Kiyama, Y., Horai, R., Sudo, K., Kakuta, S., Iwakura, Y., Iino, M., Kano, M., Manabe, T., Yamamoto, T., 2007. Involvement of protein-tyrosine phosphatase PTPMEG in motor learning and cerebellar long-term depression. Eur. J. Neurosci. 26, 2269–2278. https://doi.org/10.1111/j.1460-9568.2007.05829.x

Kiraly, D.D., Lemtiri-Chlieh, F., Levine, E.S., Mains, R.E., Eipper, B.A., 2011. Kalirin Binds the NR2B Subunit of the NMDA Receptor, Altering Its Synaptic Localization and Function. J. Neurosci. 31, 12554–12565. https://doi.org/10.1523/JNEUROSCI.3143-11.2011

Koch, I., Schwarz, H., Beuchle, D., Goellner, B., Langegger, M., Aberle, H., 2008. Drosophila Ankyrin 2 Is Required for Synaptic Stability. Neuron 58, 210–222. https://doi.org/10.1016/j.neuron.2008.03.019

Koefoed, P., Hansen, T.V.O., Woldbye, D.P.D., Werge, T., Mors, O., Hansen, T., Jakobsen, K.D., Nordentoft, M., Wang, A., Bolwig, T.G., Rehfeld, J.F., 2009. An intron 1 polymorphism in the cholecystokinin-A receptor gene associated with schizophrenia in males. Acta Psychiatr. Scand. 120, 281–287. https://doi.org/10.1111/j.1600-0447.2009.01400.x

Kohda, K., Kakegawa, W., Matsuda, S., Yamamoto, T., Hirano, H., Yuzaki, M., 2013. The 2 glutamate receptor gates long-term depression by coordinating interactions between two AMPA receptor phosphorylation sites. Proc. Natl. Acad. Sci. 110, E948–E957. https://doi.org/10.1073/pnas.1218380110

Koide, T., Banno, M., Aleksic, B., Yamashita, S., Kikuchi, T., Kohmura, K., Adachi, Y., Kawano, N., Kushima, I., Nakamura, Y., Okada, T., Ikeda, M., Ohi, K., Yasuda, Y., Hashimoto, R., Inada, T., Ujike, H., Iidaka, T., Suzuki, M., Takeda, M., Iwata, N., Ozaki, N., 2012. Common variants in MAGI2 gene are associated with increased risk for cognitive impairment in Schizophrenic patients. PLoS One 7, e36836. https://doi.org/10.1371/journal.pone.0036836

Korber, C., Werner, M., Kott, S., Ma, Z.-L., Hollmann, M., 2007. The Transmembrane AMPA Receptor Regulatory Protein 4 Is a More Effective Modulator of AMPA Receptor Function than Stargazin ( 2). J. Neurosci. 27, 8442–8447. https://doi.org/10.1523/JNEUROSCI.0424-07.2007

Krapivinsky, G., Medina, I., Krapivinsky, L., Gapon, S., Clapham, D.E., 2004. SynGAP-MUPP1-CaMKII synaptic complexes regulate p38 MAP kinase activity and NMDA receptor- dependent synaptic AMPA receptor potentiation. Neuron 43, 563–574. https://doi.org/10.1016/j.neuron.2004.08.003

Krolewski, R.C., Packard, A., Jang, W., Wildner, H., Schwob, J.E., 2012. Ascl1 (Mash1) knockout perturbs differentiation of nonneuronal cells in olfactory epithelium. PLoS One 7, e51737. https://doi.org/10.1371/journal.pone.0051737

LaRese, T.P., Yan, Y., Eipper, B.A., Mains, R.E., 2017. Using Kalirin conditional knockout mice to distinguish its role in dopamine receptor mediated behaviors. BMC Neurosci. 18, 45. https://doi.org/10.1186/s12868-017-0363-2

Lee, J., Chung, C., Ha, S., Lee, D., Kim, D.-Y., Kim, H., Kim, E., 2015. Shank3-mutant mice lacking exon 9 show altered excitation/inhibition balance, enhanced rearing, and spatial memory deficit. Front. Cell. Neurosci. 9, 94. https://doi.org/10.3389/fncel.2015.00094

Lee, J.R., 2015. Protein tyrosine phosphatase PTPRT as a regulator of synaptic formation and neuronal development. BMB Rep. https://doi.org/10.5483/BMBRep.2015.48.5.037

Lei, S., Lu, W.Y., Xiong, Z.G., Orser, B.A., Valenzuela, C.F., MacDonald, J.F., 1999. Platelet-derived growth factor receptor-induced feed-forward inhibition of excitatory transmission between hippocampal pyramidal neurons. J. Biol. Chem. 274, 30617–30623. https://doi.org/10.1074/jbc.274.43.30617

Levinson, D.F., Duan, J., Oh, S., Wang, K., Sanders, A.R., Shi, J., Zhang, N., Mowry, B.J., Olincy, A., Amin, F., Cloninger, C.R., Silverman, J.M., Buccola, N.G., Byerley, W.F., Black, D.W., Kendler, K.S., Freedman, R., Dudbridge, F., Pe'er, I., Hakonarson, H., Bergen, S.E., Fanous, A.H., Holmans, P.A., Gejman, P. V, 2011. Copy number variants in schizophrenia: Confirmation of five previous finding sand new evidence for 3q29 microdeletions and VIPR2 duplications. Am. J. Psychiatry 168, 302–316. https://doi.org/10.1176/appi.ajp.2010.10060876

Li, M., Niu, F., Zhu, X., Wu, X., Shen, N., Peng, X., Liu, Y., 2015. PRRT2 mutant leads to dysfunction of glutamate signaling. Int. J. Mol. Sci. 16, 9134–9151. https://doi.org/10.3390/ijms16059134

Lian, G., Dettenhofer, M., Lu, J., Downing, M., Chenn, A., Wong, T., Sheen, V., 2016. Filamin A- and formin 2-dependent endocytosis regulates proliferation via the canonical Wnt pathway. Development 143, 4509–4520. https://doi.org/10.1242/dev.139295

Lim, S.H., Kwon, S.K., Lee, M.K., Moon, J., Jeong, D.G., Park, E., Kim, S.J., Park, B.C., Lee, S.C., Ryu, S.E., Yu, D.Y., Chung, B.H., Kim, E., Myung, P.K., Lee, J.R., 2009. Synapse formation regulated by protein tyrosine phosphatase receptor T through interaction with cell adhesion molecules and Fyn. EMBO J. 28, 3564–3578. https://doi.org/10.1038/emboj.2009.289

Lin, R., Karpa, K., Kabbani, N., Goldman-Rakic, P., Levenson, R., 2001. Dopamine D2 and D3 receptors are linked to the actin cytoskeleton via interaction with filamin A. Proc Natl Acad Sci U S A 98, 5258–5263. https://doi.org/10.1073/pnas.011538198

Lipina, T. V, Prasad, T., Yokomaku, D., Luo, L., Connor, S.A., Kawabe, H., Wang, Y.T., Brose, N., Roder, J.C., Craig, A.M., 2016. Cognitive Deficits in Calsyntenin-2-deficient Mice Associated with Reduced GABAergic Transmission. Neuropsychopharmacology 41, 802–10. https://doi.org/10.1038/npp.2015.206

Liu, Y.-H., Tsai, J.-W., Chen, J.-L., Yang, W.-S., Chang, P.-C., Cheng, P.-L., Turner, D.L., Yanagawa, Y., Wang, T.-W., Yu, J.-Y., 2017. Ascl1 promotes tangential migration and confines migratory routes by induction of Ephb2 in the telencephalon. Sci. Rep. 7, 42895. https://doi.org/10.1038/srep42895

Liu, Y.-T., Nian, F.-S., Chou, W.-J., Tai, C.-Y., Kwan, S.-Y., Chen, C., Kuo, P.-W., Lin, P.-

H., Chen, C.-Y., Huang, C.-W., Lee, Y.-C., Soong, B.-W., Tsai, J.-W., 2016. *PRRT2* mutations lead to neuronal dysfunction and neurodevelopmental defects. Oncotarget 7, 39184–39196. https://doi.org/10.18632/oncotarget.9258

Llorente-Folch, I., Sahún, I., Contreras, L., Casarejos, M.J., Grau, J.M., Saheki, T., Mena, M.A., Satrústegui, J., Dierssen, M., Pardo, B., 2013. AGC1-malate aspartate shuttle activity is critical for dopamine handling in the nigrostriatal pathway. J. Neurochem. 124, 347–362. https://doi.org/10.1111/jnc.12096

Ma, X.-M., Kiraly, D.D., Gaier, E.D., Wang, Y., Kim, E.-J., Levine, E.S., Eipper, B.A., Mains, R.E., 2008. Kalirin-7 Is Required for Synaptic Structure and Function. J. Neurosci. 28, 12368–12382. https://doi.org/10.1523/JNEUROSCI.4269-08.2008

Maher, B.J., LoTurco, J.J., 2012. Disrupted-in-schizophrenia (DISC1) functions presynaptically at glutamatergic synapses. PLoS One 7, e34053. https://doi.org/10.1371/journal.pone.0034053

Mandela, P., Ma, X.-M., 2012. Kalirin, a Key Player in Synapse Formation, Is Implicated in Human Diseases. Neural Plast. 2012, 1–9. https://doi.org/10.1155/2012/728161

Mansilla, A., Chaves-Sanjuan, A., Campillo, N.E., Semelidou, O., Martínez-González, L., Infantes, L., González-Rubio, J.M., Gil, C., Conde, S., Skoulakis, E.M.C., Ferrús, A., Martínez, A., Sánchez-Barrena, M.J., 2017. Interference of the complex between NCS-1 and Ric8a with phenothiazines regulates synaptic function and is an approach for fragile X syndrome. Proc. Natl. Acad. Sci. 114, E999–E1008. https://doi.org/10.1073/pnas.1611089114

Maximov, A., Shin, O.H., Liu, X., Südhof, T.C., 2007. Synaptotagmin-12, a synaptic vesicle phosphoprotein that modulates spontaneous neurotransmitter release. J. Cell Biol. 176, 113–124. https://doi.org/10.1083/jcb.200607021

McNeely, K.C., Cupp, T.D., Little, J.N., Janisch, K.M., Shrestha, A., Dwyer, N.D., 2017. Mutation of Kinesin-6 Kif20b causes defects in cortical neuron polarization and morphogenesis. Neural Dev. 12, 5. https://doi.org/10.1186/s13064-017-0082-5

Milstein, A.D., Zhou, W., Karimzadegan, S., Bredt, D.S., Nicoll, R.A., 2007. TARP Subtypes Differentially and Dose-Dependently Control Synaptic AMPA Receptor Gating. Neuron 55, 905–918. https://doi.org/10.1016/j.neuron.2007.08.022

Moens, L.N., Ceulemans, S., Alaerts, M., Van Den Bossche, M.J.A., Lenaerts, A.S., De Zutter, S., Norrback, K.F., Adolfsson, R., Del-Favero, J., 2010. PCM1 and schizophrenia: A replication study in the northern Swedish population. Am. J. Med. Genet. Part B Neuropsychiatr. Genet. 153, 1240–1243. https://doi.org/10.1002/ajmg.b.31088

Mohamad, O., Song, M., Wei, L., Yu, S.P., 2013. Regulatory roles of the NMDA receptor GluN3A subunit in locomotion, Pain perception and cognitive functions in adult mice. J. Physiol. 591, 149–168. https://doi.org/10.1113/jphysiol.2012.239251

Mosca, T.J., Luginbuhl, D.J., Wang, I.E., Luo, L., 2017. Presynaptic LRP4 promotes synapse number and function of excitatory CNS neurons. Elife 6, e27347. https://doi.org/10.7554/eLife.27347

Nadler, J.V., Bray, S.D., Evenson, D.A., 1992. Autoradiographic localization of proline uptake in excitatory hippocampal pathways. Hippocampus 2, 269–278. https://doi.org/10.1002/hipo.450020306

Namekata, K., Harada, C., Taya, C., Guo, X., Kimura, H., Parada, L.F., Harada, T., 2010. Dock3 induces axonal outgrowth by stimulating membrane recruitment of the WAVE complex. Proc. Natl. Acad. Sci. 107, 7586–7591. https://doi.org/10.1073/pnas.0914514107

Namekata, K., Watanabe, H., Guo, X., Kittaka, D., Kawamura, K., Kimura, A., Harada, C., Harada, T., 2012. Dock3 regulates BDNF-TrkB signaling for neurite outgrowth by forming a ternary complex with Elmo and RhoG. Genes to Cells 17, 688–697. https://doi.org/10.1111/j.1365-2443.2012.01616.x

Nanou, E., Kyriakatos, A., Bhattacharjee, A., Kaczmarek, L.K., Paratcha, G., El Manira, A., 2008. Na+-mediated coupling between AMPA receptors and KNa channels shapes synaptic transmission. Proc. Natl. Acad. Sci. U. S. A. 105, 20941–20946. https://doi.org/10.1073/pnas.0806403106

Naruhashi, K., Kadomatsu, K., Igakura, T., Fan, Q.-W., Kuno, N., Muramatsu, H., Miyauchi, T., Hasegawa, T., Itoh, A., Muramatsu, T., Nabeshima, T., 1997. Abnormalities of Sensory and Memory Functions in Mice LackingBsgGene. Biochem. Biophys. Res. Commun. 236, 733–737. https://doi.org/10.1006/bbrc.1997.6993

Niesler, B., Flohr, T., Nöthen, M.M., Fischer, C., Rietschel, M., Franzek, E., Albus, M., Propping, P., Rappold, G.A., 2001a. Association between the 5′ utr variant c178t of the serotonin receptor gene htr3a and bipolar affective disorder. Pharmacogenetics 11, 471–475. https://doi.org/10.1097/00008571-200108000-00002

Niesler, B., Weiss, B., Fischer, C., Nöthen, M.M., Propping, P., Bondy, B., Rietschel, M., Maier, W., Albus, M., Franzek, E., Rappold, G.A., 2001b. Serotonin receptor gene HTR3A variants in schizophrenic and bipolar affective patients. Pharmacogenetics 11, 21–27. https://doi.org/10.1097/00008571-200102000-00003

Nishimura, T., Harashima, S. ichi, Yafang, H., Notkins, A.L., 2010. IA-2 modulates dopamine secretion in PC12 cells. Mol. Cell. Endocrinol. 315, 81–86. https://doi.org/10.1016/j.mce.2009.09.023

Nishimura, T., Kubosaki, A., Ito, Y., Notkins, A.L., 2009. Disturbances in the secretion of neurotransmitters in IA-2/IA-2β null mice: Changes in behavior, learning and lifespan. Neuroscience 159, 427–437. https://doi.org/10.1016/j.neuroscience.2009.01.022

Nunokawa, A., Watanabe, Y., Kaneko, N., Sugai, T., Yazaki, S., Arinami, T., Ujike, H., Inada, T., Iwata, N., Kunugi, H., Sasaki, T., Itokawa, M., Ozaki, N., Hashimoto, R., Someya, T., 2010. The dopamine D3 receptor (DRD3) gene and risk of schizophrenia: Case-control studies and an updated meta-analysis. Schizophr. Res. 116, 61–67. https://doi.org/10.1016/j.schres.2009.10.016

O'Keeffe, G.C., Tyers, P., Aarsland, D., Dalley, J.W., Barker, R.A., Caldwell, M.A., 2009. Dopamine-induced proliferation of adult neural precursor cells in the mammalian subventricular zone is mediated through EGF. Proc. Natl. Acad. Sci. 106, 8754–8759. https://doi.org/10.1073/pnas.0803955106

Ori-McKenney, K.M., Vallee, R.B., 2011. Neuronal migration defects in the loa dynein mutant mouse. Neural Dev. 6, 26. https://doi.org/10.1186/1749-8104-6-26

Ota, V.K., Bellucco, F.T., Gadelha, A., Santoro, M.L., Noto, C., Christofolini, D.M., Assunção, I.B., Yamada, K.M., Ribeiro-dos-Santos, Â.K., Santos, S., Mari, J.J., Smith, M.A.C., Melaragno, M.I., Bressan, R.A., Sato, J.R., Jackowski, A.P., Belangero, S.I., 2014. PRODH polymorphisms, cortical volumes and thickness in schizophrenia. PLoS One 9, e87686. https://doi.org/10.1371/journal.pone.0087686

Ou, X.-M., Chen, K., Shih, J.C., 2006. Monoamine oxidase A and repressor R1 are involved in apoptotic signaling pathway. Proc. Natl. Acad. Sci. U. S. A. 103, 10923–8. https://doi.org/10.1073/pnas.0601515103

Panaccione, I., Napoletano, F., Forte, A., Kotzalidis, G., Casale, A., Rapinesi, C., Brugnoli, C., Serata, D., Caccia, F., Cuomo, I., Ambrosi, E., Simonetti, A., Savoja, V., Chiara, L., Danese, E., Manfredi, G., Janiri, D., Motolese, M., Nicoletti, F., Girardi, P., Sani, G., 2013. Neurodevelopment in Schizophrenia: The Role of the Wnt Pathways. Curr. Neuropharmacol. 11, 535–558. https://doi.org/10.2174/1570159X113119990037

Parker, L.L., Backstrom, J.R., Sanders-Bush, E., Shieh, B.H., 2003. Agonist-induced phosphorylation of the serotonin 5-HT2C receptor regulates its interaction with multiple PDZ protein 1. J. Biol. Chem. 278, 21576–21583. https://doi.org/10.1074/jbc.M210973200

Paterlini, M., Zakharenko, S.S., Lai, W.S., Qin, J., Zhang, H., Mukai, J., Westphal, K.G.C., Olivier, B., Sulzer, D., Pavlidis, P., Siegelbaum, S.A., Karayiorgou, M., Gogos, J.A., 2005. Transcriptional and behavioral interaction between 22q11.2 orthologs modulates schizophrenia-related phenotypes in mice. Nat. Neurosci. 8, 1586–1594. https://doi.org/10.1038/nn1562

Pattyn, A., Guillemot, F., Brunet, J.F., 2006. Delays in neuronal differentiation in Mash1/Ascl1 mutants. Dev. Biol. 295, 67–75. https://doi.org/10.1016/j.ydbio.2006.03.008

Pavlopoulos, E., Trifilieff, P., Chevaleyre, V., Fioriti, L., Zairis, S., Pagano, A., Malleret, G., Kandel, E.R., 2011. Neuralized1 activates CPEB3: A function for nonproteolytic ubiquitin in synaptic plasticity and memory storage. Cell 147, 1369–1383. https://doi.org/10.1016/j.cell.2011.09.056

Paylor, R., Glaser, B., Mupo, A., Ataliotis, P., Spencer, C., Sobotka, A., Sparks, C., Choi, C.-H., Oghalai, J., Curran, S., Murphy, K.C., Monks, S., Williams, N., O'Donovan, M.C., Owen, M.J., Scambler, P.J., Lindsay, E., 2006. Tbx1 haploinsufficiency is linked to behavioral disorders in mice and humans: Implications for 22q11 deletion syndrome. Proc. Natl. Acad. Sci. 103, 7729–7734. https://doi.org/10.1073/pnas.0600206103

Peça, J., Feliciano, C., Ting, J.T., Wang, W., Wells, M.F., Venkatraman, T.N., Lascola, C.D., Fu, Z., Feng, G., 2011. Shank3 mutant mice display autistic-like behaviours and striatal dysfunction. Nature 472, 437–442. https://doi.org/10.1038/nature09965

Peykov, S., Berkel, S., Schoen, M., Weiss, K., Degenhardt, F., Strohmaier, J., Weiss, B., Proepper, C., Schratt, G., Nöthen, M.M., Boeckers, T.M., Rietschel, M., Rappold, G.A., 2015. Identification and functional characterization of rare SHANK2 variants in schizophrenia. Mol. Psychiatry 20, 1489–1498. https://doi.org/10.1038/mp.2014.172

Pohlkamp, T., Durakoglugil, M., Lane-Donovan, C., Xian, X., Johnson, E.B., Hammer, R.E., Herz, J., 2015. Lrp4 domains differentially regulate limb/brain development and synaptic plasticity. PLoS One 10, e0116701. https://doi.org/10.1371/journal.pone.0116701

Proulx, É., Young, E.J., Osborne, L.R., Lambe, E.K., 2010. Enhanced prefrontal serotonin 5-HT1A currents in a mouse model of Williams-Beuren syndrome with low innate anxiety. J. Neurodev. Disord. 2, 99–108. https://doi.org/10.1007/s11689-010-9044-5

Raber, J., Wong, D., Buttini, M., Orth, M., Bellosta, S., Pitas, R.E., Mahley, R.W., Mucke, L., 2002. Isoform-specific effects of human apolipoprotein E on brain function revealed in ApoE knockout mice: Increased susceptibility of females. Proc. Natl. Acad. Sci. 95, 10914–10919. https://doi.org/10.1073/pnas.95.18.10914

Ramírez, V.T., Ramos-Fernández, E., Henríquez, J.P., Lorenzo, A., Inestrosa, N.C., 2016. Wnt-5a/frizzled9 receptor signaling through the Gαo-Gβγ complex regulates dendritic spine formation. J. Biol. Chem. 291, 19092–19107. https://doi.org/10.1074/jbc.M116.722132

Ranneva, S.V., Pavlov, K.S., Gromova, A.V., Amstislavskaya, T.G., Lipina, T.V., 2017. Features of emotional and social behavioral phenotypes of calsyntenin2 knockout mice. Behav. Brain Res. 332, 343–354. https://doi.org/10.1016/j.bbr.2017.06.029

Rao, N.R., Abad, C., Perez, I.C., Srivastava, A.K., Young, J.I., Walz, K., 2017. Rai1 Haploinsufficiency Is Associated with Social Abnormalities in Mice. Biology (Basel). 6, 25. https://doi.org/http://dx.doi.org/10.3390/biology6020025

Raynaud, F., Janossy, A., Dahl, J., Bertaso, F., Perroy, J., Varrault, A., Vidal, M., Worley, P.F., Boeckers, T.M., Bockaert, J., Marin, P., Fagni, L., Homburger, V., 2013. Shank3-Rich2 interaction regulates AMPA receptor recycling and synaptic long-term potentiation. J. Neurosci. 33, 9699–9715. https://doi.org/10.1523/JNEUROSCI.2725-12.2013

Reith, R.M., McKenna, J., Wu, H., Hashmi, S.S., Cho, S.H., Dash, P.K., Gambello, M.J., 2013. Loss of Tsc2 in Purkinje cells is associated with autistic-like behavior in a mouse model of tuberous sclerosis complex. Neurobiol. Dis. 51, 93–103. https://doi.org/10.1016/j.nbd.2012.10.014

Rejeb, I., Ben Jemaa, L., Abaied, L., Kraoua, L., Saillour, Y., Maazoul, F., Chelly, J., Chaabouni, H., 2011. A novel frame shift mutation in the PQBP1 gene identified in a Tunisian family with X-linked mental retardation. Eur. J. Med. Genet. 54, 241–246. https://doi.org/10.1016/j.ejmg.2011.01.010

Robertshaw, E., Matsumoto, K., Lumsden, A., Kiecker, C., 2013. Irx3 and Pax6 establish differential competence for Shh-mediated induction of GABAergic and glutamatergic neurons of the thalamus. Proc. Natl. Acad. Sci. 110, E3919–E3926. https://doi.org/10.1073/pnas.1304311110

Romero-Pozuelo, J., Dason, J.S., Mansilla, A., Banos-Mateos, S., Sardina, J.L., Chaves-Sanjuan, A., Jurado-Gomez, J., Santana, E., Atwood, H.L., Hernandez-Hernandez, A., Sanchez-Barrena, M.-J., Ferrus, A., 2014. The guanine-exchange factor Ric8a binds to the Ca2+ sensor NCS-1 to regulate synapse number and neurotransmitter release. J. Cell Sci. 127, 4246–4259. https://doi.org/10.1242/jcs.152603

Ru, W., Peng, Y., Zhong, L., Tang, S.J., 2012. A role of the mammalian target of rapamycin (mTOR) in glutamate-induced down-regulation of tuberous sclerosis complex proteins 2 (TSC2). J. Mol. Neurosci. 47, 340–345. https://doi.org/10.1007/s12031-012-9753-1

Ruat, M., Traiffort, E., 2013. Roles of the calcium sensing receptor in the central nervous system. Best Pract. Res. Clin. Endocrinol. Metab. 27, 429–442. https://doi.org/10.1016/j.beem.2013.03.001

Ruisu, K., Kask, K., Meier, R., Saare, M., Raid, R., Veraksitš, A., Karis, A., Tõnissoo, T., Pooga, M., 2013. Ablation of RIC8A Function in Mouse Neurons Leads to a Severe Neuromuscular Phenotype and Postnatal Death. PLoS One 8, e74031. https://doi.org/10.1371/journal.pone.0074031

Sakai, K., Shoji, H., Kohno, T., Miyakawa, T., Hattori, M., 2016. Mice that lack the C-Terminal region of Reelin exhibit behavioral abnormalities related to neuropsychiatric disorders. Sci. Rep. 6, 28636. https://doi.org/10.1038/srep28636

Sanjuan, J., Toirac, I., González, J.C., Leal, C., Moltó, M.D., Nájera, C., De Frutos, R., 2004. A possible association between the CCK-AR gene and persistent auditory hallucinations in schizophrenia. Eur. Psychiatry 19, 349–353. https://doi.org/10.1016/j.eurpsy.2004.06.015

Sansom, S.N., Griffiths, D.S., Faedo, A., Kleinjan, D.J., Ruan, Y., Smith, J., Van Heyningen, V., Rubenstein, J.L., Livesey, F.J., 2009. The level of the transcription factor Pax6 is essential for controlling the balance between neural stem cell self-renewal and neurogenesis. PLoS Genet. 5, e1000511. https://doi.org/10.1371/journal.pgen.1000511

Sasaki, J., Kofuji, S., Itoh, R., Momiyama, T., Takayama, K., Murakami, H., Chida, S., Tsuya, Y., Takasuga, S., Eguchi, S., Asanuma, K., Horie, Y., Miura, K., Davies, E.M., Mitchell, C., Yamazaki, M., Hirai, H., Takenawa, T., Suzuki, A., Sasaki, T., 2010. The PtdIns(3,4)P2 phosphatase INPP4A is a suppressor of excitotoxic neuronal death. Nature 465, 497–501. https://doi.org/10.1038/nature09023

Sato, A., 2016. mTOR, a Potential Target to Treat Autism Spectrum Disorder. CNS Neurol. Disord. Drug Targets 15, 533–43. https://doi.org/10.2174/1871527315666160413120638

Sato, A., Kasai, S., Kobayashi, T., Takamatsu, Y., Hino, O., Ikeda, K., Mizuguchi, M., 2012. Rapamycin reverses impaired social interaction in mouse models of tuberous sclerosis complex. Nat. Commun. 3, 1292. https://doi.org/10.1038/ncomms2295

Schmeisser, M.J., Ey, E., Wegener, S., Bockmann, J., Stempel, A.V., Kuebler, A., Janssen, A.L., Udvardi, P.T., Shiban, E., Spilker, C., Balschun, D., Skryabin, B. V., Dieck, S. tom, Smalla, K.H., Montag, D., Leblond, C.S., Faure, P., Torquet, N., Le Sourd, A.M., Toro, R., Grabrucker, A.M., Shoichet, S.A., Schmitz, D., Kreutz, M.R., Bourgeron, T., Gundelfinger, E.D., Boeckers, T.M., 2012. Autistic-like behaviours and hyperactivity in mice lacking ProSAP1/Shank2. Nature 486, 256–260. https://doi.org/10.1038/nature11015

Scholz, R., Berberich, S., Rathgeber, L., Kolleker, A., Köhr, G., Kornau, H.C., 2010. AMPA Receptor Signaling through BRAG2 and Arf6 Critical for Long-Term Synaptic Depression. Neuron 66, 768–780. https://doi.org/10.1016/j.neuron.2010.05.003

Šekeljić, V., Andjus, P.R., 2012. Tenascin-C and its functions in neuronal plasticity. Int. J. Biochem. Cell Biol. 44, 825–829. https://doi.org/10.1016/j.biocel.2012.02.014

Setou, M., Nakagawa, T., Seog, D.H., Hirokawa, N., 2000. Kinesin superfamily motor protein KIF17 and mLin-10 in NMDA receptor- containing vesicle transport. Science (80-. ). 288, 1796–1802. https://doi.org/10.1126/science.288.5472.1796

Shen, E.Y., Jiang, Y., Mao, W., Futai, K., Hock, H., Akbarian, S., 2015. Cognition and mood-related behaviors in L3mbtl1 null mutant mice. PLoS One 10, e0121252. https://doi.org/10.1371/journal.pone.0121252

Shigetomi, E., Tong, X., Kwan, K.Y., Corey, D.P., Khakh, B.S., 2012. TRPA1 channels regulate astrocyte resting calcium and inhibitory synapse efficacy through GAT-3. Nat. Neurosci. 15, 70–80. https://doi.org/10.1038/nn.3000

Shin, H., Wyszynski, M., Huh, K.H., Valtschanoff, J.G., Lee, J.R., Ko, J., Streuli, M., Weinberg, R.J., Sheng, M., Kim, E., 2003. Association of the kinesin motor KIF1A with the multimodular protein liprin-α. J. Biol. Chem. 278, 11393–11401. https://doi.org/10.1074/jbc.M211874200

Singh, U., Kumar, S., Shelkar, G.P., Yadav, M., Kokare, D.M., Goswami, C., Lechan, R.M., Singru, P.S., 2016. Transient receptor potential vanilloid 3 (TRPV3) in the ventral tegmental area of rat: Role in modulation of the mesolimbic-dopamine reward pathway. Neuropharmacology 110, 198–210. https://doi.org/10.1016/j.neuropharm.2016.04.012

Skinbjerg, M., Ariano, M.A., Thorsell, A., Heilig, M., Halldin, C., Innis, R.B., Sibley, D.R., 2009. Arrestin3 mediates D(2) dopamine receptor internalization. Synapse 63, 621–624. https://doi.org/10.1002/syn.20636

Smith, K.R., Kopeikina, K.J., Fawcett-Patel, J.M., Leaderbrand, K., Gao, R., Schürmann, B., Myczek, K., Radulovic, J., Swanson, G.T., Penzes, P., 2014. Psychiatric Risk Factor ANK3/Ankyrin-G Nanodomains Regulate the Structure and Function of Glutamatergic Synapses. Neuron 84, 399–415. https://doi.org/10.1016/j.neuron.2014.10.010

Smith, M.J., Pozo, K., Brickley, K., Stephenson, F.A., 2006. Mapping the GRIF-1 binding domain of the kinesin, KIF5C, substantiates a role for GRIF-1 as an adaptor protein in the anterograde trafficking of cargoes. J. Biol. Chem. 281, 27216–27228. https://doi.org/10.1074/jbc.M600522200

Soler-Llavina, G.J., Arstikaitis, P., Morishita, W., Ahmad, M., Südhof, T.C., Malenka, R.C., 2013. Leucine-rich repeat transmembrane proteins are essential for maintenance of long-term potentiation. Neuron 79, 439–446. https://doi.org/10.1016/j.neuron.2013.06.007

Sousa, K.M., Carlos Villaescusa, J., Cajanek, L., Ondr, J.K., Castelo-Branco, G., Hofstra, W., Bryja, V., Palmberg, C., Bergman, T., Wainwright, B., Lang, R.A., Arenas, E., 2010. Wnt2 regulates progenitor proliferation in the developing ventral midbrain. J. Biol. Chem. 285, 7246–7253. https://doi.org/10.1074/jbc.M109.079822

Spence, E.F., Kanak, D.J., Carlson, B.R., Soderling, S.H., 2016. The Arp2/3 Complex Is Essential for Distinct Stages of Spine Synapse Maturation, Including Synapse Unsilencing. J. Neurosci. 36, 9696–9709. https://doi.org/10.1523/JNEUROSCI.0876-16.2016

Spoelgen, R., 2005. LRP2/megalin is required for patterning of the ventral telencephalon. Development 132, 405–414. https://doi.org/10.1242/dev.01580

Spuch, C., Ortolano, S., Navarro, C., 2012. LRP-1 and LRP-2 receptors function in the membrane neuron. Trafficking mechanisms and proteolytic processing in alzheimer's disease. Front. Physiol. https://doi.org/10.3389/fphys.2012.00269

Stamenkovic, V., Milenkovic, I., Galjak, N., Todorovic, V., Andjus, P., 2017. Enriched environment alters the behavioral profile of tenascin-C deficient mice. Behav. Brain Res. 331, 241–253. https://doi.org/10.1016/j.bbr.2017.05.047

Stottmann, R.W., Tran, P. V., Turbe-Doan, A., Beier, D.R., 2009. Ttc21b is required to restrict sonic hedgehog activity in the developing mouse forebrain. Dev. Biol. 335, 166–178. https://doi.org/10.1016/j.ydbio.2009.08.023

Studniarczyk, D., Coombs, I., Cull-Candy, S.G., Farrant, M., 2013. TARP γ-7 selectively enhances synaptic expression of calcium-permeable AMPARs. Nat. Neurosci. 16, 1266–1274. https://doi.org/10.1038/nn.3473

Sun, B., Bang, S.I., Jin, Y.H., 2009. Transient receptor potential A1 increase glutamate release on brain stem neurons. Neuroreport 20, 1002–1006. https://doi.org/10.1097/WNR.0b013e32832d2219

Sunmonu, N.A., Li, K., Guo, Q., Li, J.Y.H., 2011. Gbx2 and Fgf8 are sequentially required for formation of the midbrain-hindbrain compartment boundary. Development 138, 725–734. https://doi.org/10.1242/dev.055665

Tamura, T., Horiuchi, D., Chen, Y.-C., Sone, M., Miyashita, T., Saitoe, M., Yoshimura, N., Chiang, A.-S., Okazawa, H., 2010. Drosophila PQBP1 Regulates Learning Acquisition at Projection Neurons in Aversive Olfactory Conditioning. J. Neurosci. 30, 14091–14101. https://doi.org/10.1523/JNEUROSCI.1319-10.2010

Telley, L., Cadilhac, C., Cioni, J.M., Saywell, V., Jahannault-Talignani, C., Huettl, R.E., Sarrailh-Faivre, C., Dayer, A., Huber, A.B., Ango, F., 2016. Dual Function of NRP1 in Axon Guidance and Subcellular Target Recognition in Cerebellum. Neuron 91, 1276–1291. https://doi.org/10.1016/j.neuron.2016.08.015

Thakar, S., Wang, L., Yu, T., Ye, M., Onishi, K., Scott, J., Qi, J., Fernandes, C., Han, X., Yates, J.R., Berg, D.K., Zou, Y., 2017. Evidence for opposing roles of Celsr3 and Vangl2 in glutamatergic synapse formation. Proc. Natl. Acad. Sci. 114, E610–E618. https://doi.org/10.1073/pnas.1612062114

Tõnissoo, T., Kõks, S., Meier, R., Raud, S., Plaas, M., Vasar, E., Karis, A., 2006. Heterozygous mice with Ric-8 mutation exhibit impaired spatial memory and decreased anxiety. Behav. Brain Res. 167, 42–48. https://doi.org/10.1016/j.bbr.2005.08.025

Trinh, M.A., Kaphzan, H., Wek, R.C., Pierre, P., Cavener, D.R., Klann, E., 2012. Brain-Specific Disruption of the eIF2α Kinase PERK Decreases ATF4 Expression and Impairs Behavioral Flexibility. Cell Rep. 1, 676–688. https://doi.org/10.1016/j.celrep.2012.04.010

Tsai, L.-C.C.L., Chan, G.C.-K., Nangle, S.N., Shimizu-Albergine, M., Jones, G.L., Storm, D.R., Beavo, J.A., Zweifel, L.S., 2012. Inactivation of Pde8b enhances memory, motor performance, and protects against age-induced motor coordination decay. Genes, Brain Behav. 11, 837–847. https://doi.org/10.1111/j.1601-183X.2012.00836.x

Tuoc, T., Dere, E., Radyushkin, K., Pham, L., Nguyen, H., Tonchev, A.B., Sun, G., Ronnenberg, A., Shi, Y., Staiger, J.F., Ehrenreich, H., Stoykova, A., 2017. Ablation of BAF170 in Developing and Postnatal Dentate Gyrus Affects Neural Stem Cell Proliferation, Differentiation, and Learning. Mol. Neurobiol. 54, 4618–4635. https://doi.org/10.1007/s12035-016-9948-5

Tuoc, T.C., Boretius, S., Sansom, S.N., Pitulescu, M.E., Frahm, J., Livesey, F.J., Stoykova, A., 2013. Chromatin Regulation by BAF170 Controls Cerebral Cortical Size and Thickness. Dev. Cell 25, 256–269. https://doi.org/10.1016/j.devcel.2013.04.005

Tuoc, T.C., Radyushkin, K., Tonchev, A.B., Pinon, M.C., Ashery-Padan, R., Molnar, Z., Davidoff, M.S., Stoykova, A., 2009. Selective Cortical Layering Abnormalities and Behavioral Deficits in Cortex-Specific Pax6 Knock-Out Mice. J. Neurosci. 29, 8335–8349. https://doi.org/10.1523/JNEUROSCI.5669-08.2009

Tzavara, E.T., Bymaster, F.P., Davis, R.J., Wade, M.R., Perry, K.W., Wess, J., McKinzie, D.L., Felder, C., Nomikos, G.G., 2004. M4 muscarinic receptors regulate the dynamics of cholinergic and dopaminergic neurotransmission: relevance to the pathophysiology and treatment of related CNS pathologies. FASEB J. 18, 1410–1412. https://doi.org/10.1096/fj.04-1575fje

Uchino, S., Wada, H., Honda, S., Hirasawa, T., Yanai, S., Nakamura, Y., Ondo, Y., Kohsaka, S., 2003. Slo2 sodium-activated K+channels bind to the PDZ domain of PSD-95. Biochem. Biophys. Res. Commun. 310, 1140–1147. https://doi.org/10.1016/j.bbrc.2003.09.133

Uetani, N., 2000. Impaired learning with enhanced hippocampal long-term potentiation in PTPdelta-deficient mice. EMBO J. 19, 2775–2785. https://doi.org/10.1093/emboj/19.12.2775

Um, J.W., Choii, G., Park, D., Kim, D., Jeon, S., Kang, H., Mori, T., Papadopoulos, T., Yoo, T., Lee, Y., Kim, E., Tabuchi, K., Ko, J., 2016. IQ Motif and SEC7 domain-containing protein 3 (IQSEC3) interacts with gephyrin to promote inhibitory synapse formation. J. Biol. Chem. 291, 10119–10130. https://doi.org/10.1074/jbc.M115.712893

Underhill, S.M., Wheeler, D.S., Amara, S.G., 2015. Differential Regulation of Two Isoforms of the Glial Glutamate Transporter EAAT2 by DLG1 and CaMKII. J. Neurosci. 35, 5260–5270. https://doi.org/10.1523/JNEUROSCI.4365-14.2015

Utsunomiya, K., Shinkai, T., De Luca, V., Hwang, R., Sakata, S., Fukunaka, Y., Chen, H.I., Ohmori, O., Nakamura, J., 2008. Genetic association between the dopamine D3 gene polymorphism (Ser9Gly) and schizophrenia in Japanese populations: Evidence from a case-control study and meta-analysis. Neurosci. Lett. 444, 161–165. https://doi.org/10.1016/j.neulet.2008.08.005

Valente, P., Castroflorio, E., Rossi, P., Fadda, M., Sterlini, B., Cervigni, R.I., Prestigio, C., Giovedì, S., Onofri, F., Mura, E., Guarnieri, F.C., Marte, A., Orlando, M., Zara, F., Fassio, A., Valtorta, F., Baldelli, P., Corradi, A., Benfenati, F., 2016. PRRT2 Is a Key Component of the Ca2+-Dependent Neurotransmitter Release Machinery. Cell Rep. 15, 117–131. https://doi.org/10.1016/j.celrep.2016.03.005

Valenzuela, C.F., Xiong, Z., MacDonald, J.F., Weiner, J.L., Frazier, C.J., Dunwiddie, T. V, Kazlauskas, A., Whiting, P.J., Harris, R.A., 1996. Platelet-derived growth factor induces a long-term inhibition of N-methyl-D-aspartate receptor function. J. Biol. Chem. 271, 16151–9.

Valtorta, F., Benfenati, F., Zara, F., Meldolesi, J., 2016. PRRT2: from Paroxysmal Disorders to Regulation of Synaptic Function. Trends Neurosci. https://doi.org/10.1016/j.tins.2016.08.005

van der Werf, I.M., Van Dam, D., Missault, S., Yalcin, B., De Deyn, P.P., Vandeweyer, G., Kooy, R.F., 2017. Behavioural characterization of AnkyrinG deficient mice, a model for ANK3 related disorders. Behav. Brain Res. 328, 218–226. https://doi.org/10.1016/j.bbr.2017.04.014

Vingtdeux, V., Chang, E.H., Frattini, S.A., Zhao, H., Chandakkar, P., Adrien, L., Strohl, J.J., Gibson, E.L., Ohmoto, M., Matsumoto, I., Huerta, P.T., Marambaud, P., 2016. CALHM1 deficiency impairs cerebral neuron activity and memory flexibility in mice. Sci. Rep. 6, 24250. https://doi.org/10.1038/srep24250

Voikar, V., Kulesskaya, N., Laakso, T., Lauren, J., Strittmatter, S.M., Airaksinen, M.S., 2013. LRRTM1-deficient mice show a rare phenotype of avoiding small enclosures-A tentative mouse model for claustrophobia-like behaviour. Behav. Brain Res. 238, 69–78. https://doi.org/10.1016/j.bbr.2012.10.013

Walch, L., 2013. Emerging role of the scaffolding protein Dlg1 in vesicle trafficking. Traffic. https://doi.org/10.1111/tra.12089

Walsh, T., McClellan, J.M., McCarthy, S.E., Addington, A.M., Pierce, S.B., Cooper, G.M., Nord, A.S., Kusenda, M., Malhotra, D., Bhandari, A., Stray, S.M., Rippey, C.F., Roccanova, P., Makarov, V., Lakshmi, B., Findling, R.L., Sikich, L., Stromberg, T., Merriman, B., Gogtay, N., Butler, P., Eckstrand, K., Noory, L., Gochman, P., Long, R., Chen, Z., Davis, S., Baker, C., Eichler, E.E., Meltzer, P.S., Nelson, S.F., Singleton, A.B., Lee, M.K., Rapoport, J.L., King, M.C., Sebat, J., 2008. Rare structural variants disrupt multiple genes in neurodevelopmental pathways in schizophrenia. Science (80-. ). 320, 539–543. https://doi.org/10.1126/science.1155174

Wang, H., Ge, G., Uchida, Y., Luu, B., Ahn, S., 2011. Gli3 Is Required for Maintenance and Fate Specification of Cortical Progenitors. J. Neurosci. 31, 6440–6448. https://doi.org/10.1523/JNEUROSCI.4892-10.2011

Wang, N., Xu, J., 2015. Functions of kinesin superfamily proteins in neuroreceptor trafficking. Biomed Res. Int. https://doi.org/10.1155/2015/639301

Wassef, A., Baker, J., Kochan, L.D., 2003. GABA and Schizophrenia: A Review of Basic Science and Clinical Studies. J. Clin. Psychopharmacol. 23, 601–640. https://doi.org/10.1097/01.jcp.0000095349.32154.a5

Webber, E., Li, L., Chin, L.-S., 2008. Hypertonia-associated protein Trak1 is a novel regulator of endosome-to-lysosome trafficking. J. Mol. Biol. 382, 638–51. https://doi.org/10.1016/j.jmb.2008.07.045

Wei, M., Jia, M., Zhang, J., Yu, L., Zhao, Y., Chen, Y., Ma, Y., Zhang, W., Shi, Y.S., Zhang, C., 2017. The Inhibitory Effect of α/β-Hydrolase Domain-Containing 6 (ABHD6) on the

Surface Targeting of GluA2- and GluA3-Containing AMPA Receptors. Front. Mol. Neurosci. 10, 55. https://doi.org/10.3389/fnmol.2017.00055

Wei, M., Zhang, J., Jia, M., Yang, C., Pan, Y., Li, S., Luo, Y., Zheng, J., Ji, J., Chen, J., Hu, X., Xiong, J., Shi, Y., Zhang, C., 2016. α/β-Hydrolase domain-containing 6 (ABHD6) negatively regulates the surface delivery and synaptic function of AMPA receptors. Proc. Natl. Acad. Sci. 113, E2695–E2704. https://doi.org/10.1073/pnas.1524589113

Weisenhaus, M., Allen, M.L., Yang, L., Lu, Y., Nichols, C.B., Su, T., Hell, J.W., Mcknight, G.S., 2010. Mutations in AKAP5 disrupt dendritic signaling complexes and lead to electrophysiological and behavioral phenotypes in mice. PLoS One 5, e10325. https://doi.org/10.1371/journal.pone.0010325

Williams, N.M., 2011. Molecular mechanisms in 22q11 deletion syndrome. Schizophr. Bull. https://doi.org/10.1093/schbul/sbr095

Williams, N.M., Glaser, B., Norton, N., Williams, H., Pierce, T., Moskvina, V., Monks, S., Del Favero, J., Goossens, D., Rujescu, D., Giegling, I., Kirov, G., Craddock, N., Murphy, K.C., O'Donovan, M.C., Owen, M.J., 2008. Strong evidence that GNB1L is associated with schizophrenia. Hum. Mol. Genet. 17, 555–566. https://doi.org/10.1093/hmg/ddm330

Wilson, S.L., Wilson, J.P., Wang, C., Wang, B., Mcconnell, S.K., 2012. Primary cilia and Gli3 activity regulate cerebral cortical size. Dev. Neurobiol. 72, 1196–1212. https://doi.org/10.1002/dneu.20985

Wojcik, S.M., Rhee, J.S., Herzog, E., Sigler, A., Jahn, R., Takamori, S., Brose, N., Rosenmund, C., 2004. An essential role for vesicular glutamate transporter 1 (VGLUT1) in postnatal development and control of quantal size. Proc. Natl. Acad. Sci. U. S. A. 101, 7158–63. https://doi.org/10.1073/pnas.0401764101

Won, H., Lee, H.R., Gee, H.Y., Mah, W., Kim, J.I., Lee, J.J.S., Ha, S., Chung, C., Jung, E.S., Cho, Y.S., Park, S.G., Lee, J.J.S., Lee, K., Kim, D., Bae, Y.C., Kaang, B.K., Lee, M.G., Kim, E., 2012. Autistic-like social behaviour in Shank2-mutant mice improved by restoring NMDA receptor function. Nature 486, 261–265. https://doi.org/10.1038/nature11208

Xie, Z., Cahill, M.E., Radulovic, J., Wang, J., Campbell, S.L., Miller, C.A., Sweatt, J.D., Penzes, P., 2011. Hippocampal phenotypes in kalirin-deficient mice. Mol. Cell. Neurosci. 46, 45–54. https://doi.org/10.1016/j.mcn.2010.08.005

Xu, C., Fu, X., Zhu, S., Liu, J.-J., 2016. Retrolinkin recruits the WAVE1 protein complex to facilitate BDNF-induced TrkB endocytosis and dendrite outgrowth. Mol. Biol. Cell 27, 3342–3356. https://doi.org/10.1091/mbc.E16-05-0326

Xu, J., Xiao, N., Xia, J., 2010. Thrombospondin 1 accelerates synaptogenesis in hippocampal neurons through neuroligin 1. Nat. Neurosci. 13, 22–24. https://doi.org/10.1038/nn.2459

Yamazaki, M., Fukaya, M., Hashimoto, K., Yamasaki, M., Tsujita, M., Itakura, M., Abe, M., Natsume, R., Takahashi, M., Kano, M., Sakimura, K., Watanabe, M., 2010. TARPs γ-2 and γ-7 are essential for AMPA receptor expression in the cerebellum. Eur. J. Neurosci. 31, 2204–2220. https://doi.org/10.1111/j.1460-9568.2010.07254.x

Yan, Z., Wei, J., Graziane, N.M., Wang, H., Zhong, P., Wang, Q., Liu, W., Hayashi-Takagi, A., Korth, C., Sawa, A., Brandon, N.J., 2014. Regulation of N-methyl-D-aspartate receptors by disrupted-in-schizophrenia- 1. Biol. Psychiatry 75, 414–424. https://doi.org/10.1016/j.biopsych.2013.06.009

Ying, G., Wu, S., Hou, R., Huang, W., Capecchi, M.R., Wu, Q., 2009. The Protocadherin Gene Celsr3 Is Required for Interneuron Migration in the Mouse Forebrain. Mol. Cell. Biol. 29, 3045–3061. https://doi.org/10.1128/MCB.00011-09

Yokoyama, T., Ohbuchi, T., Saito, T., Sudo, Y., Fujihara, H., Minami, K., Nagatomo, T., Uezono, Y., Ueta, Y., 2011. Allyl isothiocyanates and cinnamaldehyde potentiate miniature excitatory postsynaptic inputs in the supraoptic nucleus in rats. Eur. J. Pharmacol. 655, 31–37. https://doi.org/10.1016/j.ejphar.2011.01.011

Yoshimura, N., Horiuchi, D., Shibata, M., Saitoe, M., Qi, M. ling, Okazawa, H., 2006. Expression of human PQBP-1 in Drosophila impairs long-term memory and induces abnormal courtship. FEBS Lett. 580, 2335–2340. https://doi.org/10.1016/j.febslet.2006.03.056

Yu, H., Wang, L., Lv, L., Ma, C., Du, B., Lu, T., Jin, C., Yan, H., Yang, Y., Li, W., Ruan, Y., Zhang, Hongyan, Zhang, Hongxing, Mi, W., Mowry, B., Ma, W., Li, K., Zhang, D., Yue, W., 2016. Genome-wide association study suggested the PTPRD polymorphisms were associated with weight gain effects of atypical antipsychotic medications. Schizophr. Bull. 42, 814–823. https://doi.org/10.1093/schbul/sbv179

Yu, X., Wang, G., Gilmore, A., Yee, A.X., Li, X., Xu, T., Smith, S.J., Chen, L., Zuo, Y., 2013. Accelerated Experience-Dependent Pruning of Cortical Synapses in Ephrin-A2 Knockout Mice. Neuron 80, 64–71. https://doi.org/10.1016/j.neuron.2013.07.014

Zhang, Y. V., Hannan, S.B., Kern, J. V., Stanchev, D.T., Koç, B., Jahn, T.R., Rasse, T.M., 2017. The KIF1A homolog Unc-104 is important for spontaneous release, postsynaptic density maturation and perisynaptic scaffold organization. Sci. Rep. 7, 38172. https://doi.org/10.1038/srep38172

Zhang, Y. V., Hannan, S.B., Stapper, Z.A., Kern, J. V., Jahn, T.R., Rasse, T.M., 2016. The Drosophila KIF1A Homolog unc-104 Is Important for Site-Specific Synapse Maturation. Front. Cell. Neurosci. 10, 207. https://doi.org/10.3389/fncel.2016.00207

Zhao, C., 2005. Hippocampal and visuospatial learning defects in mice with a deletion of frizzled 9, a gene in the Williams syndrome deletion interval. Development 132, 2917–2927. https://doi.org/10.1242/dev.01871

Zhao, X.F., Kohen, R., Parent, R., Duan, Y., Fisher, G.L., Korn, M.J., Ji, L., Wan, G., Jin, J., Püschel, A.W., Dolan, D.F., Parent, J.M., Corfas, G., Murphy, G.G., Giger, R.J., 2018. PlexinA2 Forward Signaling through Rap1 GTPases Regulates Dentate Gyrus Development and Schizophrenia-like Behaviors. Cell Rep. 22, 456–470. https://doi.org/10.1016/j.celrep.2017.12.044

Zheng, C., Fu, Q., Shen, Y., Xu, Q., 2012. Investigation of allelic heterogeneity of the CCK-A receptor gene in paranoid schizophrenia. Am. J. Med. Genet. Part B Neuropsychiatr. Genet. 159B, 741–747. https://doi.org/10.1002/ajmg.b.32079

Zhou, W.J., Xu, N., Kong, L., Sun, S.C., Xu, X.F., Jia, M.Z., Wang, Y., Chen, Z.Y., 2016.

The antidepressant roles of Wnt2 and Wnt3 in stress-induced depression-like behaviors. Transl. Psychiatry 6, e892. https://doi.org/10.1038/tp.2016.122

Zhou, Z., Hu, Z.Z., Zhang, L., Hu, Z.Z., Liu, H., Liu, Z., Du, J., Zhao, J., Zhou, L., Xia, K., Tang, B., Shen, L., 2016. Identification of RELN variation p.Thr3192Ser in a Chinese family with schizophrenia. Sci. Rep. 6, 24327. https://doi.org/10.1038/srep24327

Zhu, S., Cordner, Z.A., Xiong, J., Chiu, C.-T., Artola, A., Zuo, Y., Nelson, A.D., Kim, T.-Y., Zaika, N., Woolums, B.M., Hess, E.J., Wang, X., Chuang, D.-M., Pletnikov, M.M., Jenkins, P.M., Tamashiro, K.L., Ross, C.A., 2017. Genetic disruption of ankyrin-G in adult mouse forebrain causes cortical synapse alteration and behavior reminiscent of bipolar disorder. Proc. Natl. Acad. Sci. 114, 201700689. https://doi.org/10.1073/pnas.1700689114

Zoubovsky, S., Oh, E.C., Cash-Padgett, T., Johnson, A.W., Hou, Z., Mori, S., Gallagher, M., Katsanis, N., Sawa, A., Jaaro-Peled, H., 2015. Neuroanatomical and behavioral deficits in mice haploinsufficient for Pericentriolar material 1 (Pcm1). Neurosci. Res. 98, 45–49. https://doi.org/10.1016/j.neures.2015.02.002
