## Supplemental Figure_1 for "Oligogenic rare variant contributions in schizophrenia and their convergence with genes harbouring *de novo* mutations in schizophrenia, autism and intellectual disability: Evidence from multiplex families"

**Supplementary Figure 1** : Pedigree of families used for whole exome sequencing
Showing prioritized variants encompassing genes and their transmission

- Gene names given for individuals with available DNA sample
- ■ ● represent schizophrenia affected individuals
- □ ○ Un affected individuals
- Circles and rectangles filled with other colors represent other forms of psychiatric disorders
- ✡ Representing samples used for WES
- **#** Representing the samples used for target capture sequencing

### Figure 1a : Family1

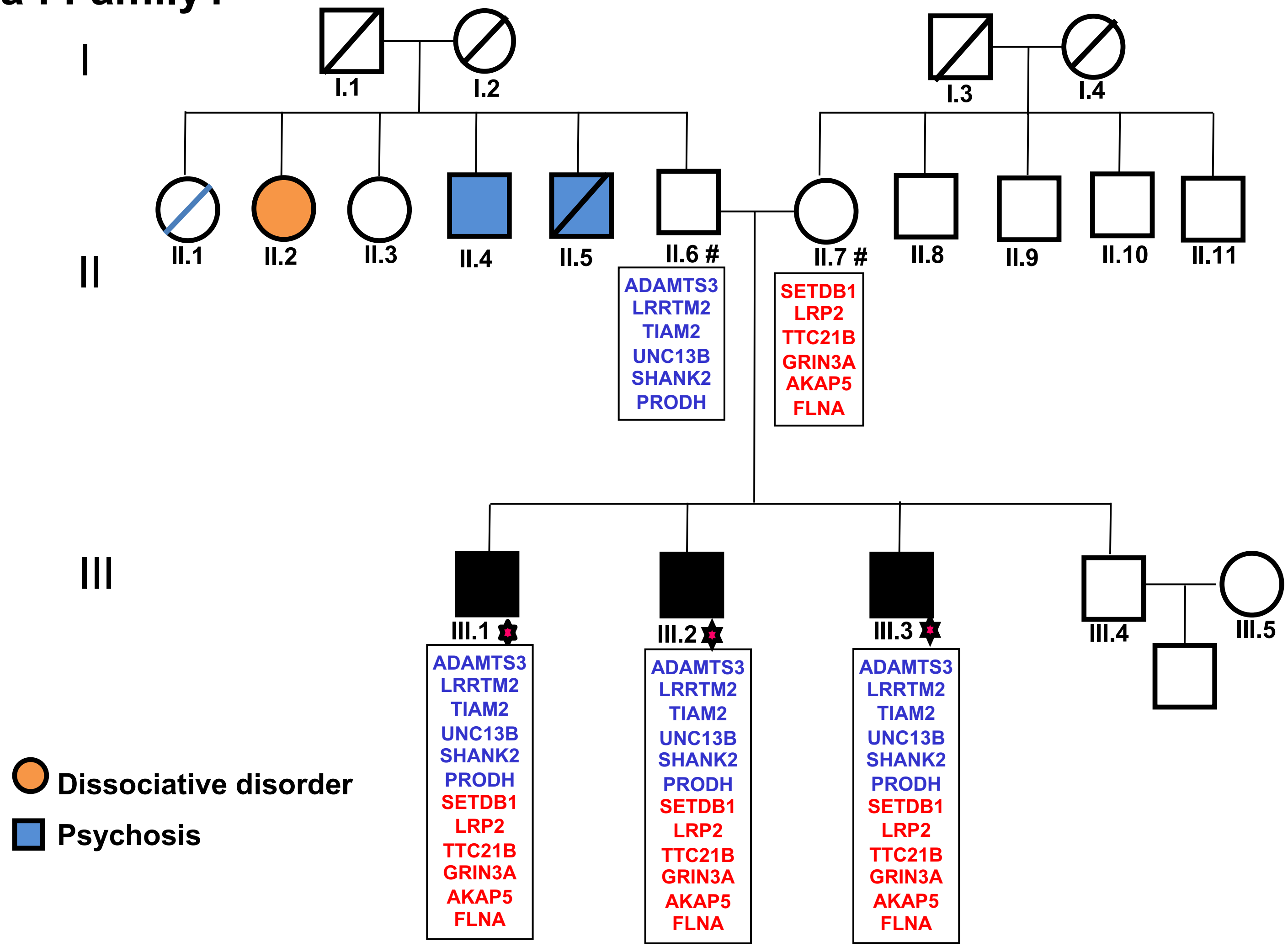

### Figure 1b : Family 2

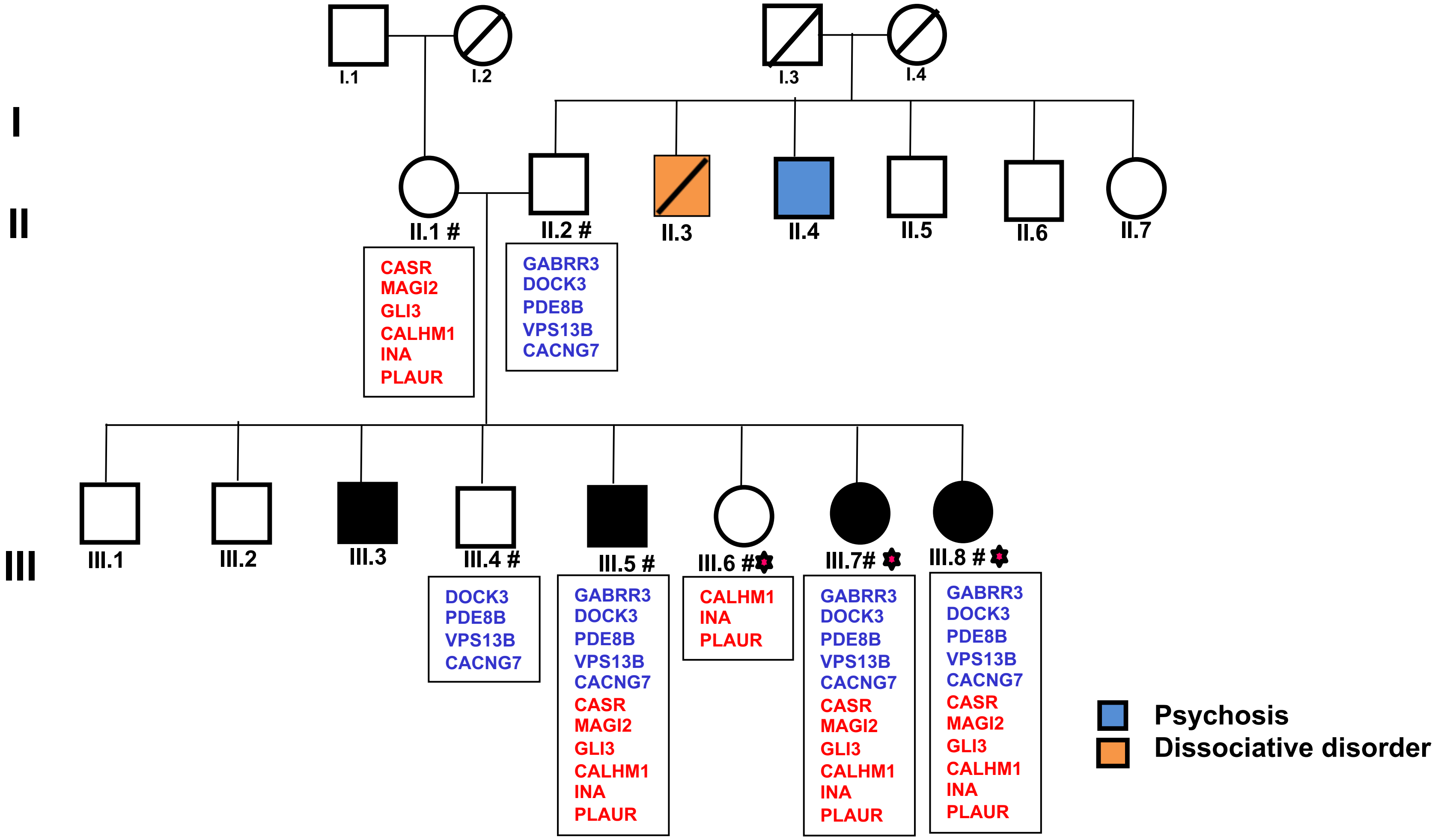

#### Figure 1c: Family 3

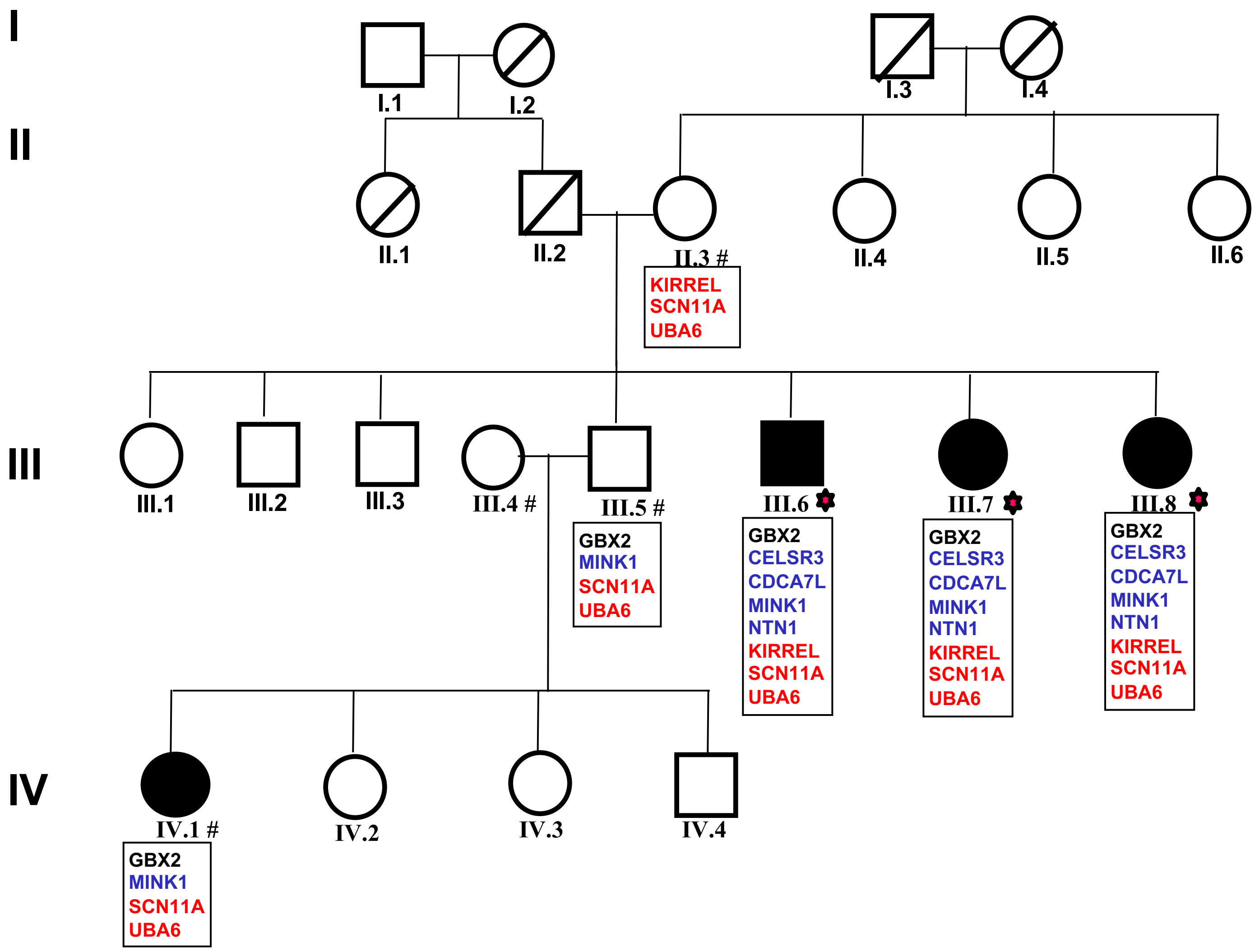

### Figure 1d: Family 4

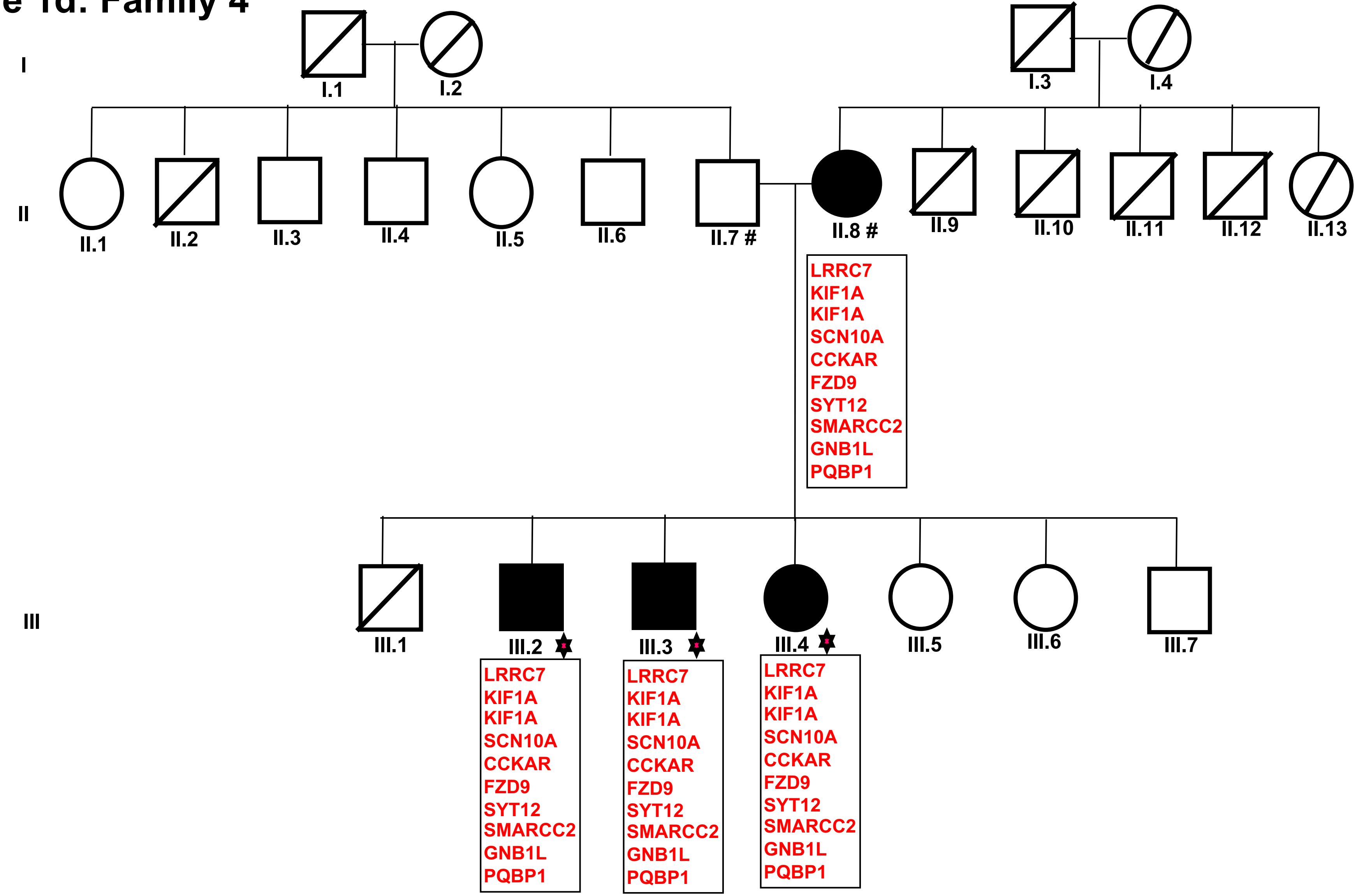

Figure 1e: Family 5

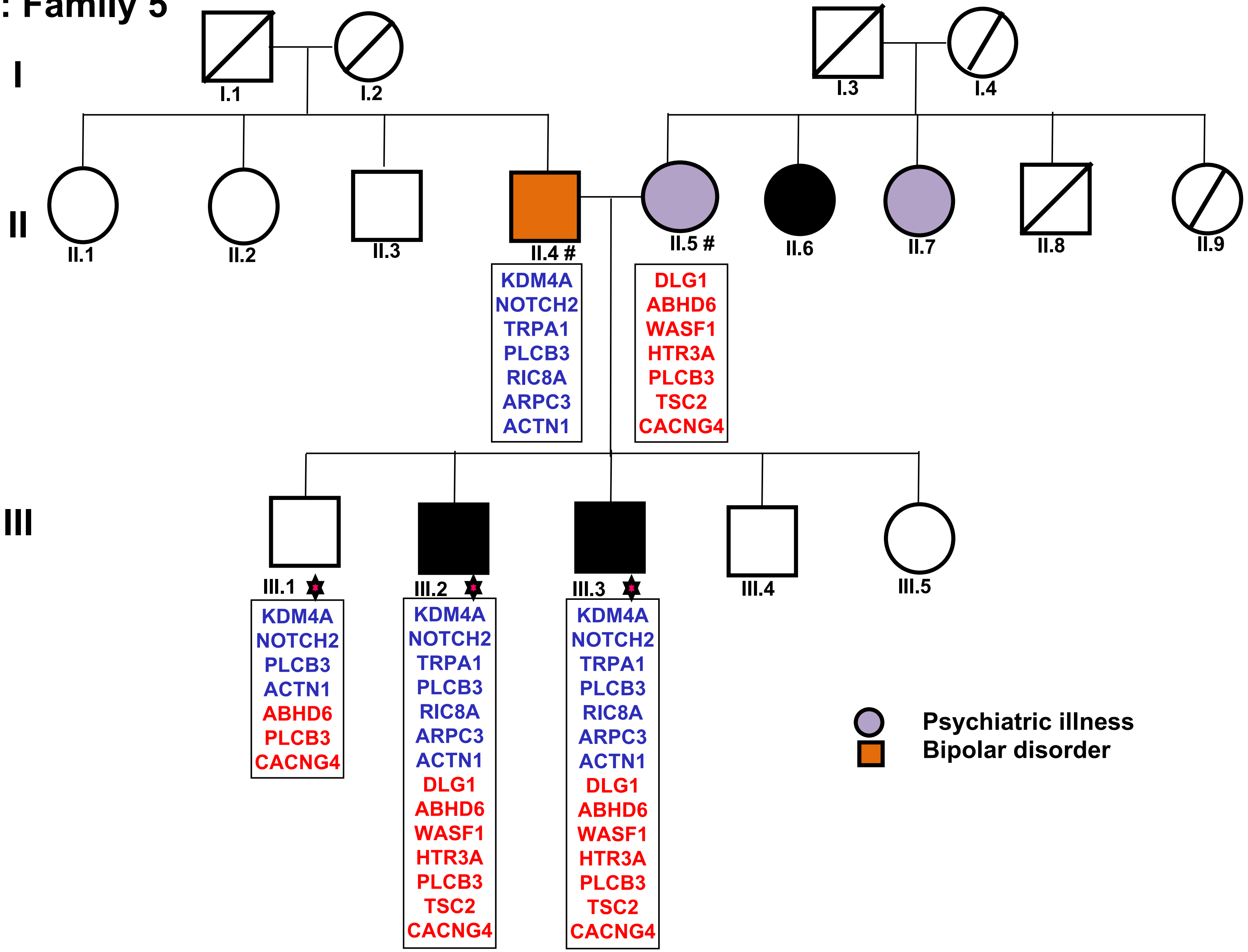

#### Figure 1f: Family 6

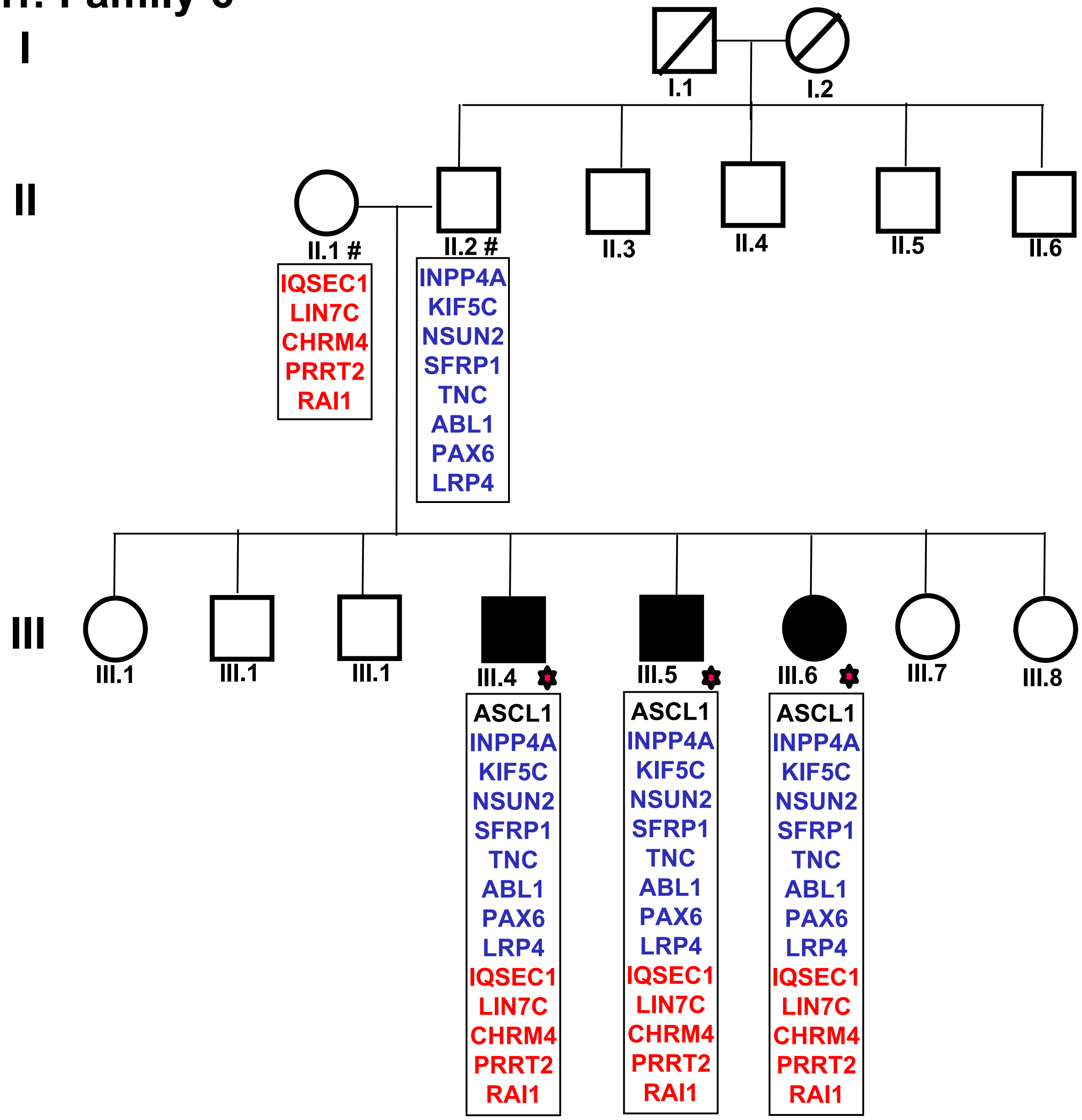

### Figure 1g: Family 7

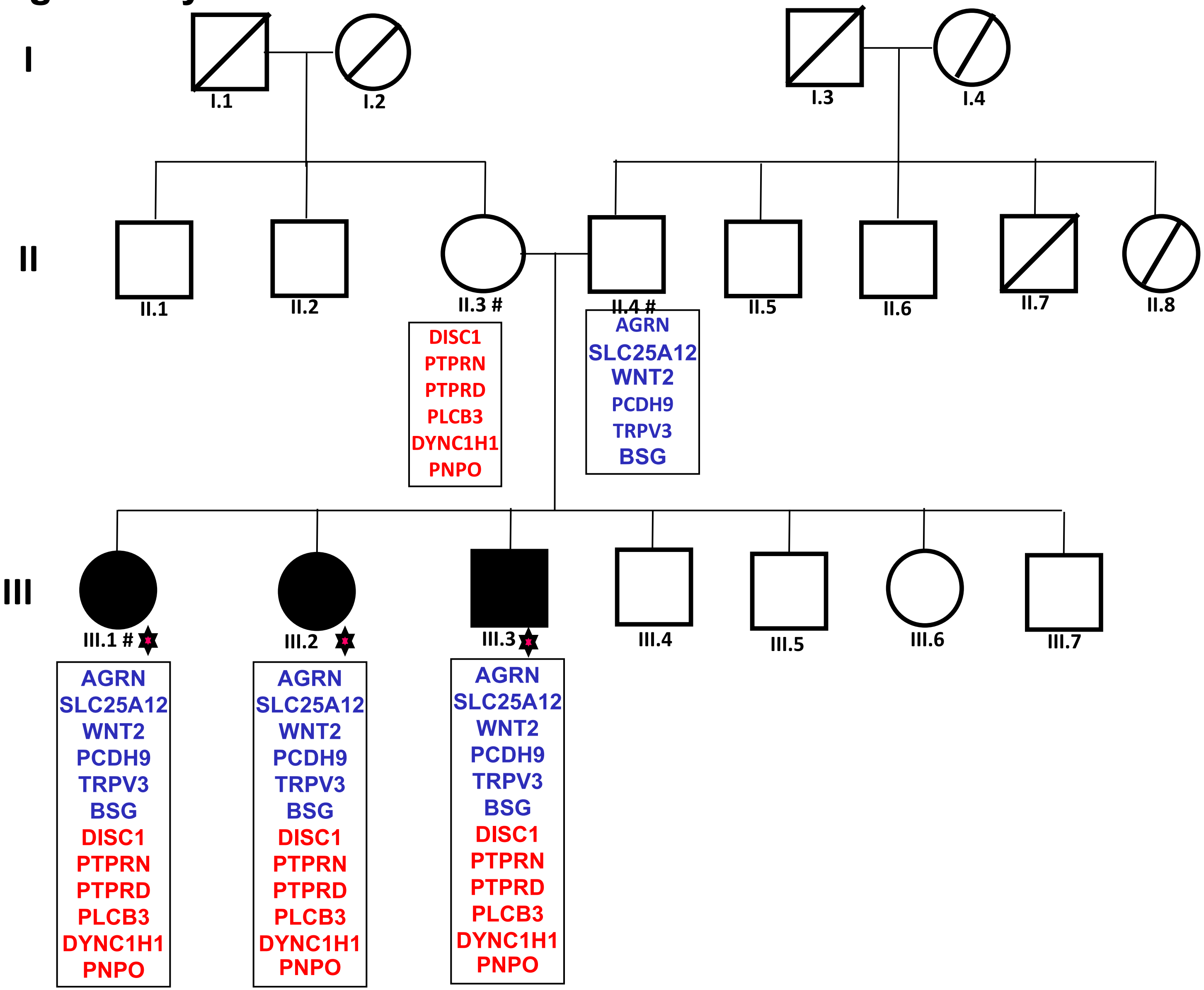

### Figure 1h: Family 8

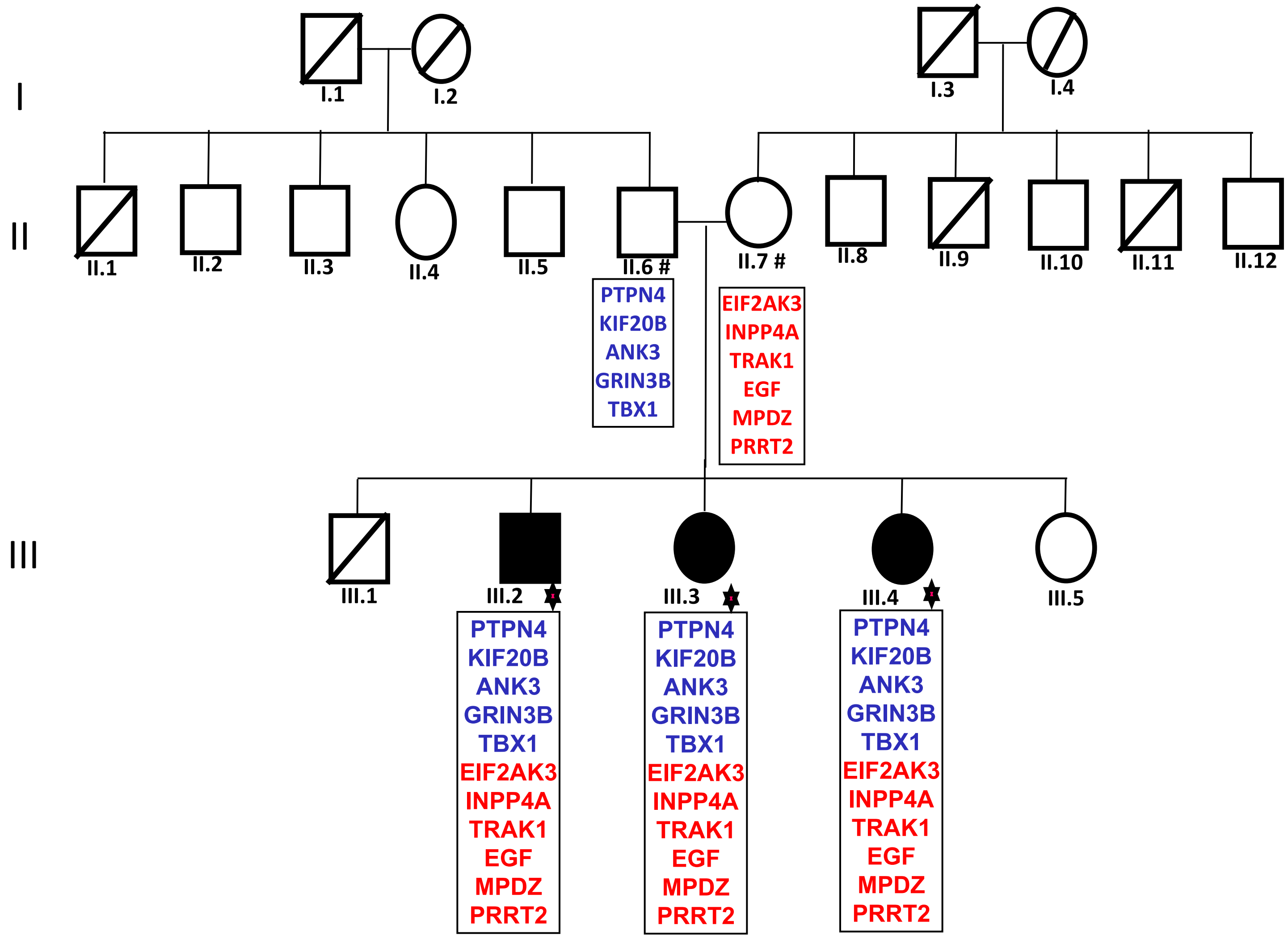

### Figure 1i: Family 9

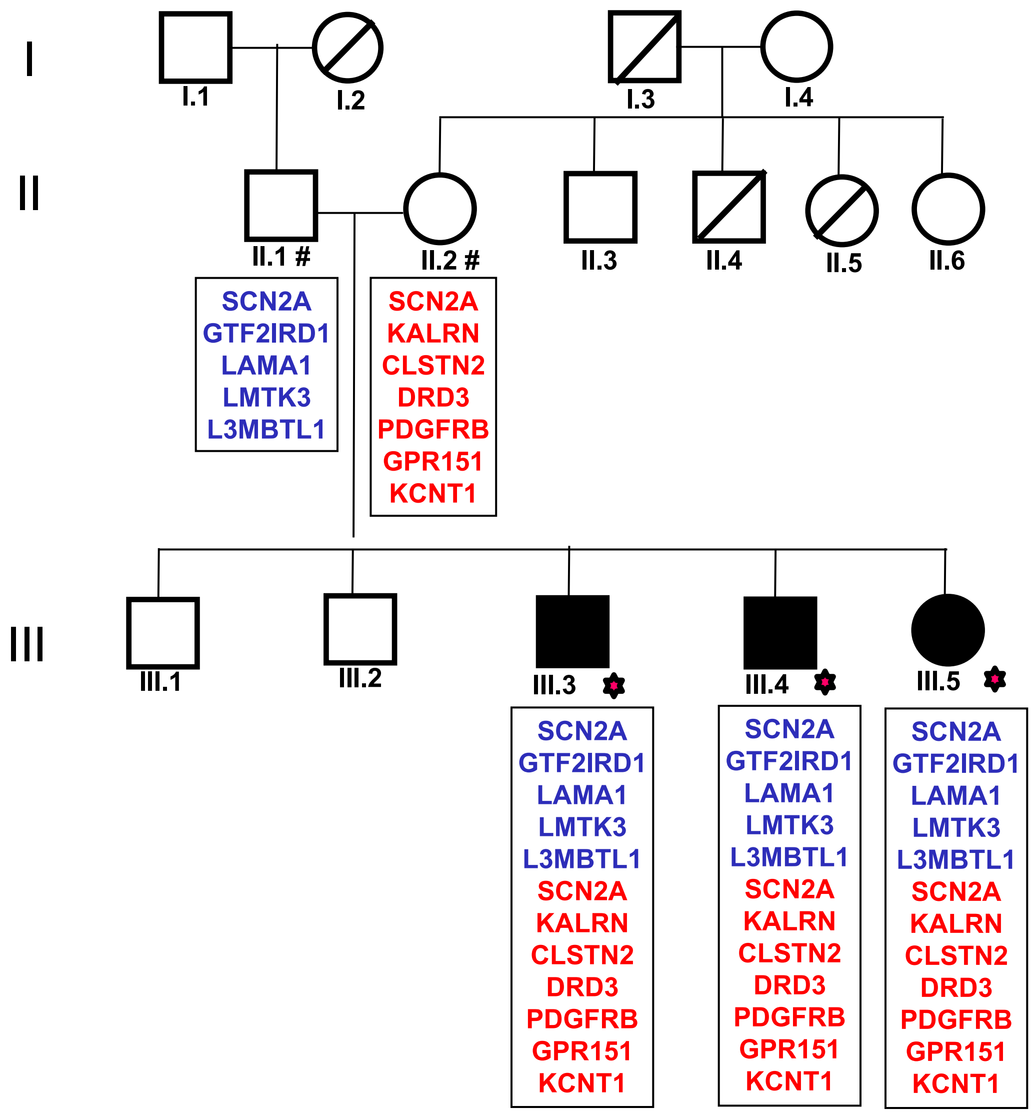

### Figure 1j: Family 10

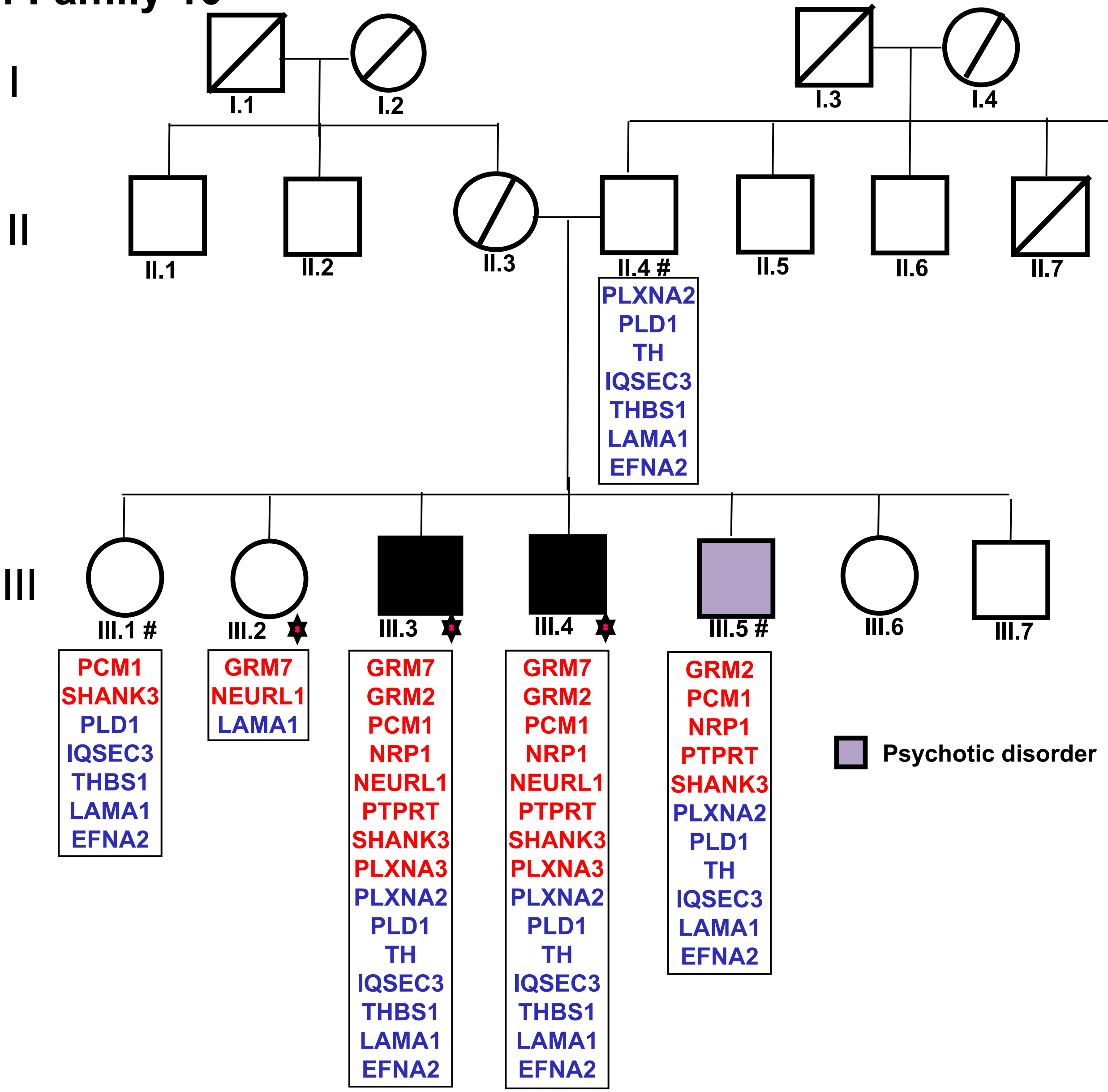

#### Figure 1k: Family 11

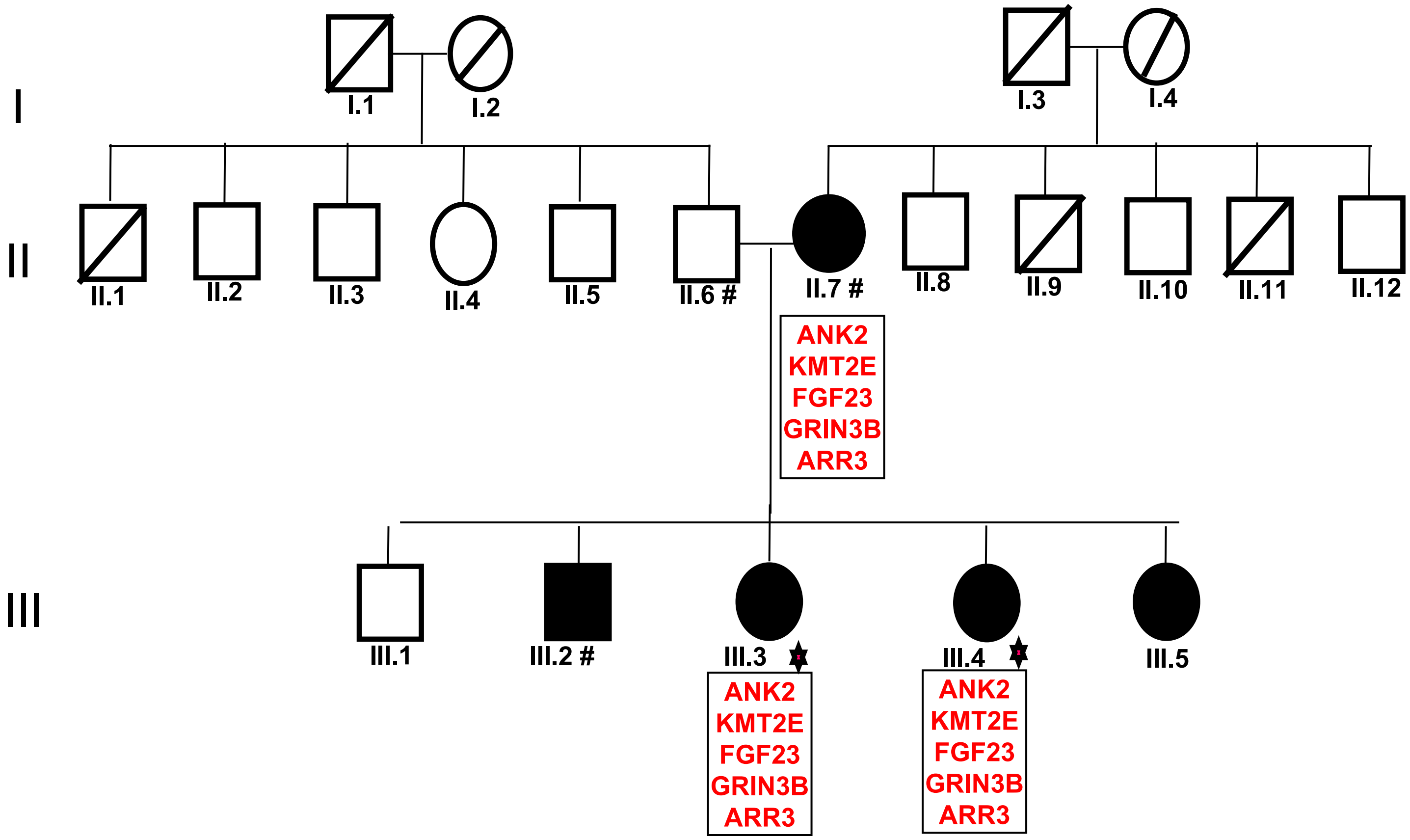
